# Interferon-γ driven differentiation of monocytes into PD-L1^+^ and MHC II^+^ macrophages and the frequency of Tim-3^+^ tumor-reactive CD8^+^ T cells within the tumor microenvironment predict a positive response to anti-PD-1-based therapy in tumor-bearing mice

**DOI:** 10.1101/2024.08.01.606242

**Authors:** Jelena Gabrilo, Sylvie Vande Velde, Coralie Henin, Sébastien Denanglaire, Abdulkader Azouz, Louis Boon, Benoit J. Van den Eynde, Muriel Moser, Stanislas Goriely, Oberdan Leo

**Author notes:** Corresponding author: Stanislas Goriely. Equal contribution (Stanislas Goriely, Oberdan Leo).

## Abstract

While immune checkpoint inhibitors have demonstrated durable responses in various cancer types, a significant proportion of patients do not exhibit favourable responses to these interventions. To uncover potential factors associated with a positive response to immunotherapy, we established a bilateral tumor model using P815 mastocytoma implanted in DBA/2 mice. In this model, only a fraction of tumor-bearing mice responds favourably to anti-PD-1 treatment, thus providing a valuable model to explore the influence of the tumor microenvironment (TME) in determining the efficacy of immune checkpoint blockade (ICB)-based immunotherapies. Moreover, this model allows for the analysis of a pretreatment tumor and inference of its treatment outcome based on the response observed in the contralateral tumor. Here, we demonstrated that tumor-reactive CD8^+^ T cell clones expressing high levels of Tim-3 were associated to a positive anti-tumor response following anti-PD-1 administration. Our study also revealed distinct differentiation dynamics in tumor-infiltrating myeloid cells in responding and non-responding mice. An IFNγ-enriched TME appeared to promote the differentiation of monocytes into PD-L1^pos^ MHC II^high^ cells in mice responding to immunotherapy. Monocytes present in the TME of non-responding mice failed to reach the same final stage of differentiation trajectory, suggesting that an altered monocyte to macrophage route may hamper the response to ICB. These insights will direct future research towards a temporal analysis of TAMs, aiming to identify factors responsible for transitions between differentiation states within the TME. This approach may potentially pave the way to novel strategies to enhance the efficacy of PD-1 blockade.

## INTRODUCTION

A large body of clinical and pre-clinical observations have illustrated the capacity of the immune system to spontaneously recognize, and eventually reject established malignancies in vivo (1). Tumors can however escape immune surveillance by displaying low immunogenicity, often a result of altered antigen presentation, or by promoting the establishment of an immunosuppressive environment opposing the development of an efficient anti-tumoral response. Recruitment of immunosuppressive cells (such as regulatory T cells (Tregs) and tumor-associated macrophages (TAMs) or neutrophils) and expression of receptors able to engage negative costimulatory molecules (such as PD-1 and CTLA-4, hence referred to as “checkpoints”) represent major mechanisms whereby tumors evade the immune response (2). These observations indicate that functional inactivation (often referred to as exhaustion) rather than clonal deletion, represents the major mechanism of peripheral tolerance to tumors. These considerations have led to the development of therapeutic strategies aimed at reestablishing the functional status of tumor infiltrating lymphocytes (TILs) by the administration of checkpoint inhibitors (such as anti-PD-1 and anti-CTLA4 antibodies, as single reagents or in combination).

Immunotherapies based on these checkpoint inhibitors have revolutionized the treatment of many cancer types over the last decade, leading to unprecedent clinical responses. Despite these undisputed therapeutic successes, only a fraction of patients responds to these immunotherapies, for reasons that are not completely understood. Current studies have identified multiple factors that can influence a positive response to checkpoint inhibitors (3), that can be broadly attributed to decreased tumor immunogenicity and/or defective lymphocyte effector function, often a consequence of the immunosuppressive tumoral microenvironment (TME).

Not surprisingly, loss of intrinsic tumor immunogenicity represents a leading cause of unresponsiveness to immunotherapy. T cell-mediated detection of neoantigens and subsequent tumor-directed cytotoxicity are critical for the successful control of tumor growth in response to immunotherapy (4–7). Tumors can however modify their antigenic profile under immune selection (a process termed immunoediting (8)), leading to a progressive loss of neoantigen expression (9) associated to resistance to anti-PD-1-based therapy (10,11). Tumors can also escape immune detection by losing or strongly reducing their antigen-processing and/or presenting capacity, a strategy that hinders expression of tumor associated antigens independently of their nature or origin (12–17).

T cell exclusion, i.e. the absence or low frequency of T cells in the TME, represents an additional, frequent cause of unresponsiveness to immunotherapy (18). These “cold” tumors, in opposition to their “hot” counterparts (i.e. tumors infiltrated by immune cells), are thought to represent most of human advanced cancers and exhibit poor response to immune checkpoint inhibitors (19,20). Although the preexisting infiltrate by tumor-reactive lymphocytes appears as a prerequisite for a successful response to immunotherapy, in situ mechanisms preventing a response to checkpoint inhibitors in “hot” tumors have been uncovered. In particular, lack of PD-L1 expression has been identified as a cause of unresponsiveness to anti-PD-1-based therapy (21) often resulting from a defective spontaneous production of the effector cytokine IFN-γ (22). Exhausted T cells represent a heterogeneous population of dysfunctional cells, of which only a subset appears to regain effector capacities upon checkpoint inhibitor administration (23–26). Although specific epigenetic modifications are probably responsible for the “reversibility” of exhausted cells, (27,28), the possible clinical translation of this concept awaits further studies.

Unresponsiveness to immunotherapy may also stem from the vast array of immunosuppressive mechanisms that operate within the TME. Tregs (29,30), and TAMs (31) represent immune-derived cells that have been shown to limit the expansion and/or effector function of antigen-specific T lymphocytes in experimental animal models. While the presence of Tregs in the tumor microenvironment has been associated to a negative prognosis in clinical studies (32,33), their specific role in opposing immune checkpoint based-immunotherapy awaits rigorous confirmation. Similarly, the impact of myeloid cells on resistance to immunotherapy in clinical setting has not been fully explored to date (34).

Although clinically relevant, the cellular and molecular data available from large cohorts of cancer patients treated with checkpoint inhibitors are often confounded by variability in tumor biology and genetics, disease stage, previous therapies, and environmental factors. A reductionist approach based on a well-defined animal model can potentially help identify critical factors that regulate the response to immunotherapy in syngeneic individuals submitted to an identical tumor burden. A set of bilateral tumor models, in which tumor-bearing mice display heterogenous responses to immunotherapy have been described (35–38). In the present study we used a P815 mastocytoma tumor model that displays a bimodal responsiveness to anti-PD-1 monotherapy. We took advantage of this observation to investigate the nature and composition of immune tumor infiltrate associated to a positive response to immunotherapy. We demonstrate herein that the presence of PD-L1-expressing myeloid cells and Tim-3^high^, tumor reactive CD8^+^ T cells in the tumor microenvironment is associated with tumor regression in response to anti-PD-1-based therapy.

## RESULTS

### A mouse model to study the influence of the tumor microenvironment in modulating the response to anti-PD-1-based immunotherapy

When implanted with the P815 mastocytoma cell line, syngeneic DBA/2 mice treated with anti-PD-1 antibodies (Figure 1A) display a bimodal response, causing a complete and persistent tumor regression in approximatively half of the mice, identified as regressors, leaving tumor growth unaltered in mice identified as progressors (Figure 1B). To identify possible roles of the TME in determining the capacity of individual mice to respond or not to this therapy, a bilateral model was developed, in which mice were inoculated with 1x10^6^ P815 cells in each flank and treated 10 days later as previously described (Figure 1C). Of note, pre-therapy tumor volume did not vary significantly between regressors and progressor throughout the study (Figure 1D). We took advantage of this model to surgically remove one of the tumors before immunotherapy to characterize the pretreatment TME, while growth of the other tumor was monitored to evaluate the progressor vs regressor state of individual mice. A series of control experiments were performed to ensure that prior anesthesia and surgery did not affect the response to anti-PD-1 Abs, and that the regressor vs progressor status was a characteristic of individual mice (i.e. tumors from individual mice behaved identically and left and right tumor volumes were significantly correlated, see Figure S1A and S1B). Finally, when tumors isolated from progressor mice were reimplanted in naïve animals, a similar dual response to immunotherapy was observed, indicating that lack of response was not due to the outgrowth of poorly immunogenic or immunoevasive tumor subclones (data not shown). Similar experiments have been performed over a period of 6 years (with n = 147 mice – 22 independent experiments), and despite some variability, the frequency of mice responding to this protocol of immunotherapy was approximatively 50% (20-80% of mice displaying a regressor phenotype, Figure S1C and S1D).

**Figure 1.**
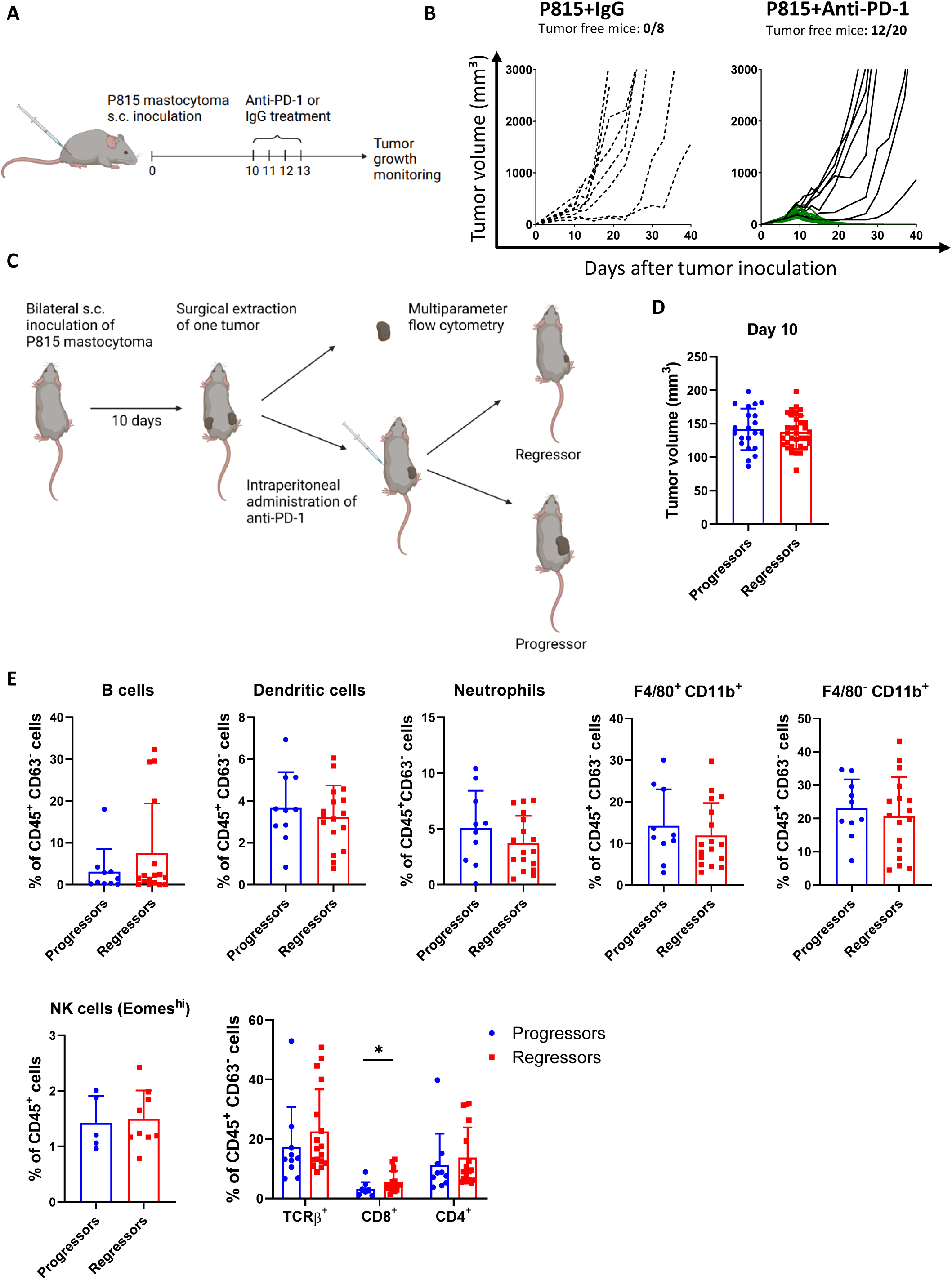
A mouse model to study the influence of the tumor microenvironment in modulating the response to anti-PD-1-based immunotherapy. 2 x 10^6^ P815 mastocytoma cells were inoculated subcutaneously (s.c.) into the right flank of DBA/2 mice and treated intraperitonealy (i.p.) with 250 µg of anti-PD-1 or isotype control at day 10, 11, 12 and 13. Tumor volume was measured 3x per week. **A)** Experimental setup. Created with BioRender.com. **B)** Growth of P815 mastocytoma tumors treated with isotype control or anti-PD-1 treatment. **C)** Schematic overview of the experimental design to study pretreatment tumor microenvironment. DBA/2 mice were inoculated s.c. with 1 x 10^6^ P815 mastocytoma cells in both flanks. Tumor infiltrating immune cells from the left flank were harvested 10 days after tumor inoculation and analyzed by flow cytometry. Mice bearing contralateral tumors were treated with 250 µg of anti-PD-1 monoclonal antibodies and tumor growth was followed. Created with BioRender.com. **D)** Initial tumor volumes in progressors and regressors at the onset of anti-PD-1 treatment. Total number of mice represents all mice used in bilateral tumor implantation experiments and subjected to surgical resection. **E)** Proportions of indicated cell populations among total immune cells at Day 10 (before anti-PD-1 treatment). Data show a pool of 2 (NK cells) or 4 independent experiments with 2-6 mice per group. Statistical comparisons were performed by using unpaired t test or Mann-Whitney test (D-E). *p < 0.05. The absence of asterisks in the graph indicates no significant difference between groups.

### Responsiveness to immunotherapy is linked to the presence of tumor-reactive CD8^+^ T cells expressing high levels of Tim-3

To evaluate the possible influence of the tumor infiltrating immune compartment on the response to immunotherapy, tumors were collected before treatment (at day 10, see Figure 1) and analyzed by flow cytometry (see Figure S2 for gating strategy). The frequency of B lymphocytes and major myeloid cell subsets was not statistically different between responding and non-responding animals. The frequency of CD8^+^ T cells among CD45^+^ tumor infiltrating cells was however increased in tumor samples of mice that subsequently responded to immunotherapy (Figure 1E). Two P815 tumor-specific antigens were previously identified (39,40). Through MHC I - tetramer staining, we evaluated the frequency of CD8^+^ TILs that reacted against the P815E antigen (a mutated form of MsrA, coding for a methionine sulfoxide reductase referred to as P1E) or against the P815A antigen (an unmutated, tumor associated antigen Trap1a, i.e. tumor rejection antigen P1A). The frequency of P1E-specific cells among CD8^+^ TILs reached 20% and did not vary between mice responding or not to anti-PD-1-based therapy (Figure 2A). In contrast, while representing a minor subset of tumor reactive lymphocytes (5 to 10%), CD8^+^ T cells reacting to P1A displayed an increased frequency in the TME of regressor mice (Figure 2B). We reached similar conclusions when we evaluated the absolute numbers of P1E-or P1A-reactive CD8^+^ T cells per gram of tumor tissue (Figure 2C and 2D). Of note, the frequency of P1A and P1E-reactive cells did not exceed 0.5 % in the spleen and lymph nodes of naïve animals, while their peripheral frequency plateaued at 1% in tumor bearing mice (Figure S3A), supporting the notion that the presence of these cells in the TME was the result of an antigen-driven process. In support of the important role of P1A-specific cells in the response to immunotherapy, a negative correlation was found between the frequency of P1A, but not P1E-reactive cells present in the TME before treatment and tumor volume following anti-PD-1 based immunotherapy (Figure 2E and 2F). This experimental model provides the opportunity to compare the phenotype of tumor reactive CD8^+^ T cells associated to distinct therapeutic consequences following immunotherapy. Both tumor-reactive subsets were found to express similar levels of PD-1 (Figure 2G), TOX and CD69 (Figure S3B). P1A-reactive CD8^+^ T cells from regressor mice were found to express high levels of markers associated to cell activation and exhaustion (such as Tim-3, CD38 and Tigit, Figure 2G and S3B). Of note, these cells were also found to express lower levels of T-bet when compared to their counterparts from progressor mice, but overall higher level compared to P1E-reactive cells, leading to a slight reduction in the TOX/T-bet ratio (Figure 2H), recently associated to cells at a stage preceding irreversible exhaustion (see (41)). Flow cytometry analysis of cell surface-associated TCRβ chains revealed a reduced expression of the TCR complex by P1E-reactive vs P1A-reactive cells in the tumor draining lymph nodes, but not in the TME (Figure 2I). Although circumstantial, this finding is compatible with the hypothesis that high avidity T lymphocytes reacting to a neoantigen may be prone to terminal exhaustion. From the functional standpoint, CD8-expressing, but not NK cells from regressor mice were found to produce higher levels of IFNγ in comparison to progressors when stimulated ex vivo by PMA/Ionomycin (Figure S4A and S4B). However, the capacity for cytokine and granzyme B production did not differ between P1E- and P1A-reactive cells (Figure S4C and S4D). Increased effector function (including IFNγ, Granzyme B and CD107 expression) was associated to the Tim-3^high^ subset, overrepresented in P1A-reactive cells from regressor mice (Figure S4E and S4F). The role of P1A-reactive cells in tumor rejection was strengthened by examining the response of DBA/2 mice to anti-PD-1-based immunotherapy when implanted with a P1A-deficient P815 subclone (Figure S5). While a majority (20 out of 30) of mice implanted with a control tumor responded to immunotherapy, this frequency was reduced to 12 out of 30 in mice inoculated with the P1A-deficient subclone. The absence of P1A-reactive clones did not result in alterations in the frequency of P1E-reactive cells (Figure S5B). Collectively these observations confirm the important role of tumor-infiltrating CD8^+^ T cells in the response to immunotherapy and suggest a possible predominant role of T cell clones expressing high levels of Tim-3 in determining tumor rejection following PD-1 blockade.

**Figure 2.**
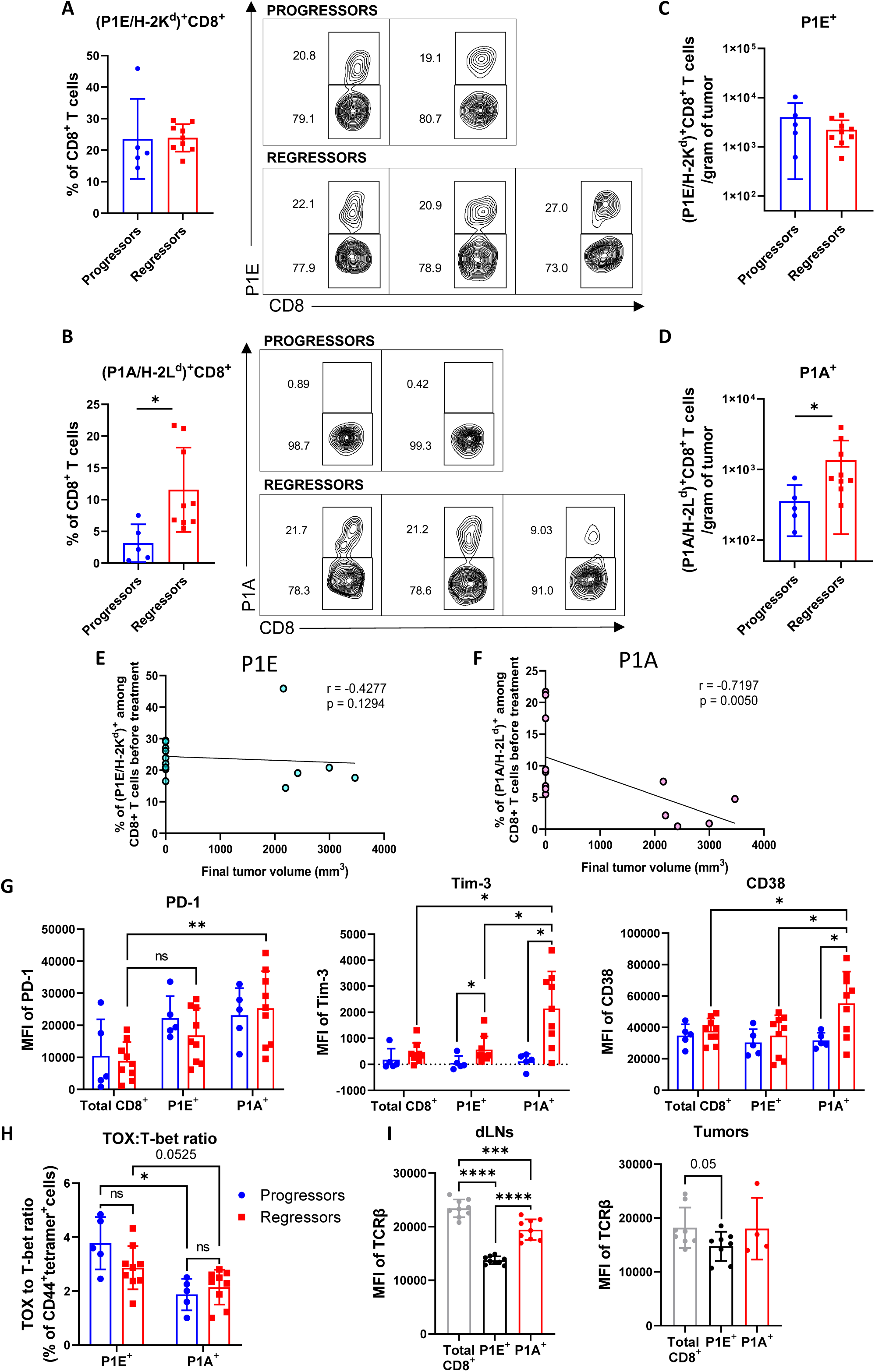
Responsiveness to immunotherapy is associated with increased infiltration of P1A-specific CD8^+^ T cells expressing high levels of Tim-3. DBA/2 mice were inoculated s.c. with 1 x 10^6^ P815 mastocytoma cells in both flanks. Tumors from one flank were surgically resected 10 days after tumor inoculation and tumor-infiltrating immune cells analyzed by flow cytometry. Mice bearing contralateral tumors were treated with 250 µg of anti-PD-1 monoclonal antibodies from day 10 to day 13 and tumor growth was followed. **A)** Proportion of P1E-specific cells among CD8^+^ T cell population and representative FACS plot. **B)** Proportion of P1A-specific cells among CD8^+^ T cell population and representative FACS plot. **C)** Number of (P1E/H-2K^d^)^+^ and **D)** (P1A/H-2L^d^)^+^ CD8^+^ T cells per gram of tumor. **E)** Correlation analysis plotting the frequency of (P1E/H-2K^d^)^+^ and **F)** (P1A/H-2L^d^)^+^ cells among CD8^+^ T cell population against final tumor volume after anti-PD-1 treatment. **G)** Median fluorescence intensity (MFI) of PD-1, Tim-3 and CD38 in total CD8^+^ population, CD44^+^ P1E- and P1A-specific CD8^+^ T cells in progressors and regressors. **H)** TOX to T-bet ratio in CD44^+^tetramer^+^ CD8^+^ T cells. **I)** MFI of TCRβ in total CD8^+^ population, CD44^+^ P1E- and P1A-specific CD8^+^ T cells in draining lymph nodes and tumors. (A-H) Data show a pool of 2 independent experiments with 2-5 mice per group or I) 1 experiment with 4-9 mice per group. Statistical comparisons were performed by using Mann-Whitney test (A-B, C, D, H) or unpaired t test (H, I), Spearman correlation (E, F) and one-way ANOVA with Tukey’s multiple comparisons test or Kruskal-Wallis test with Dunn’s multiple comparisons test (G). *p < 0.05, **p < 0.01. The absence of asterisks in the graph indicates no significant difference between groups.

### Mice responding to immunotherapy display increased PD-L1 expression by myeloid cells

The role of myeloid cells, and especially macrophages, in modulating immune control of tumor growth has been well established (31). Despite the lack of obvious correlation between the frequency of myeloid cells and the responder status of mice, we next undertook a detailed analysis of TME-associated myeloid cells. Sorted, CD11b and/or CD11c expressing cells extracted from progressor (n = 6) and regressor (n = 9) mice were submitted to single-cell RNA sequencing (scRNA-seq). Unsupervised clustering of 14,585 analysed cells led to the identification of 13 distinct single-cell clusters which were subsequently visualized using Uniform Manifold Approximation and Projection (UMAP). We detected two major meta-clusters: one comprised of monocytes and macrophages (M_1 - M_9) and another consisting of dendritic cells (DC_1 - DC_4) (Figure 3A and 3B). Dendritic cells were characterized by their elevated expression of MHC class II genes (*H2-Aa*, *H2-Ab1*, *H2-Eb1*, *H2-DMb1*, *H2-DMb2*) and *Ciita*, except for cluster DC_3, which was recognized as plasmacytoid DC expressing *Siglech* and *Ccr9* genes. DC_1 cluster expressed *Cd209a*, *Clec10a* and *Mgl2* and resembled to the T-bet^-^ cDC2 cluster previously identified by Brown and colleagues (42). DC_2 most likely represented mature dendritic cells (*Cd80*, *Cd83*, *Cd40*) enriched in immunoregulatory molecules such as *Cd274* and *Pdcd1lg2*. Cells within this cluster also expressed *Fscn1*, *Ccr7*, *Ccl22* and were analogous to mreg DC, as previously defined by Maier and colleagues (43). DC_4, identified as *Xcr1*^+^*Clec9a*^+^*Itgae*^+^ (CD103), also expressed *Batf3* and *Irf8*, known to be important for the development of type 1 conventional DCs, cDC1. Additionally, these cells were found to express genes associated with cell proliferation and cell cycle (*Mki67* and *Top2a*), suggesting that they may represent proliferating cDC1 cells. No difference in gene expression was found between the DC subsets extracted from progressor and regressor mice. Notably, a significant increase in the frequency of T-bet^-^ cDC2 cells (DC_1) was observed in the progressor group (Figure 3C and D).

**Figure 3.**
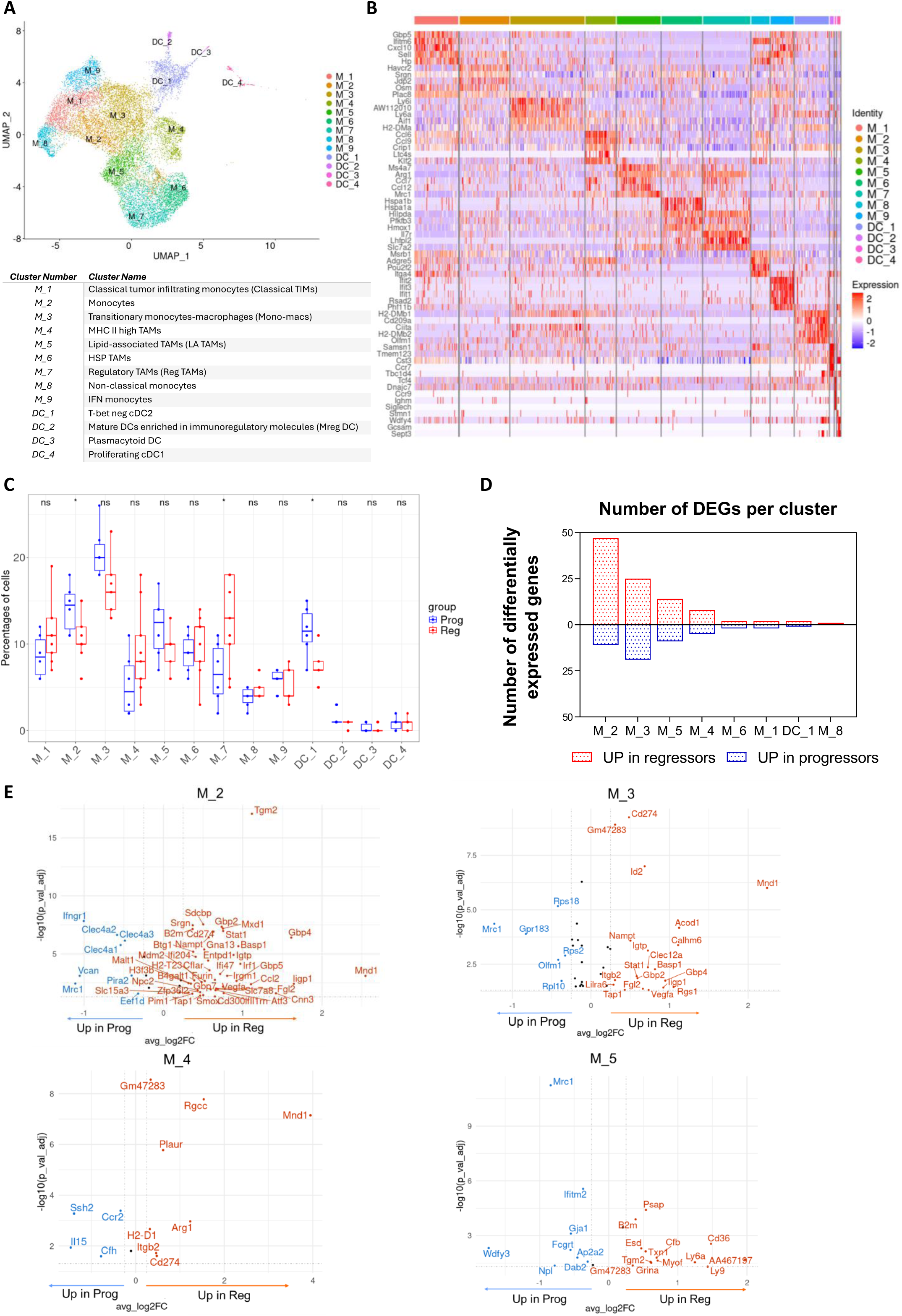
Transcriptional profiling reveals distinct gene signatures in progressors and regressors and associates baseline expression of myeloid PD-L1 to a positive therapeutic outcome in response to anti-PD-1 treatment. **A)** Uniform manifold approximation and projection (UMAP) of scRNA-seq data from CD11b^+^CD11c^+^Ly6G^-^ myeloid cells from progressors (n=6) and regressors (n=9). Colors indicate scRNA-seq clusters. **B)** Heat map of the top 5 marker genes for each cluster. **C)** Frequencies of each cluster within groups of progressors and regressors. **D)** Number of differentially expressed genes (DEGs) in indicated clusters. **E)** Volcano plots of DEGs between progressors and regressors for indicated clusters. Genes with upregulated expression in regressors are represented in red, while genes with upregulated expression in progressors are represented in blue. Data show a pool of 3 independent experiments. Statistical significance was assessed by Wilcoxon-Mann-Whitney test (C).

Clusters M_1 to M_9 represented monocytes and TAMs. Broadly, cells were divided into monocytic clusters and TAMs based on the expression levels of markers such as *Ly6c2*, *Ccr2*, *Lyz2*, *H2-Aa*, *H2-Ab1*, and *H2-Eb1*, which exhibited expression across multiple populations. Clusters M_1, M_2, M_3, M_8 and M_9 comprised cells expressing monocyte markers *Ly6c2* and *Lyz2*. M_1 and M_2 shared the expression of classical monocytic signature genes *Vcan* and *Sell* which indicated their potential recent migration from the bloodstream. However, M_1 differed from M_2 by high levels of *Mgst1*, *Cxcl10* and *Ifitm6* expression therefore showing a resemblance to the classical tumor infiltrating monocytes (TIMs). These TIMs were thought to represent the initial phase of monocyte-to-macrophage differentiation. On the other hand, M_2 exhibited significant expression of *Havcr2* and *Osm*. Cluster M_3 cells acquired the expression of MHC class II genes (*H2-Ab1*, *H2-Eb1*, *H2-Aa*) and other genes regulated by IFNγ (*Stat1*, *Cxcl9*, *Ly6a*). This cluster shared signature with both monocytes and macrophages, and was thus annotated as Mono-macs, reflecting an ongoing process of monocyte-to-macrophage differentiation. The M_3 and M_9 subsets shared an overlapping gene signature which includes *Stat1* and *Cxcl9*. However, M_9 cells expressed additional interferon (IFN)-associated genes, including *Ifit1/2/3*, *Isg15*, *Rsad2*, that were not observed in M_3. Due to the absence of MHC class II gene expression, these cells were thought to be IFN-regulated monocytes, likely possessing a pro-inflammatory function. M_8 was distinct as it contained a small portion of cells expressing transcripts associated with the non-classical monocytes (*Ace*, *Ear2*, *Adgre4*, *Ceacam1*, *Nr4a1*).

M_4 to M_7 clusters displayed features of more differentiated cells, with the majority of them characterized by the expression of immunoregulatory genes such as *Arg1* or *Trem2*, leading us to consider them as TAMs. TREM2 has been shown to play an important role in regulating immunosuppressive activity of myeloid cells (44). These clusters mainly displayed lower expression of MHC, with the exception of M_4 which was the only TAM cluster still maintaining high MHC class II genes expression. Additionally, the M_4 cluster displayed high expression of genes associated with complement activation, such as *C1qa*, *C1qb*, and *C1qc* as well as *Ccl6* and *Ccl9*. This cluster retained low expression of *Ly6c2*, suggesting that these cells may have potentially differentiated from the mono-macro cluster, following the same differentiation trajectory. M_5 possessed a signature of lipid-related genes that includes *Apoe*, *Ctsb*, *Ctsd* and *Ctsl*, along with *Ms4a7*, *Ccl7* and *Ccl12*. This cluster demonstrated the highest expression levels of *Mrc1* and was assumed to be linked to the lipid metabolism (see Figure 3B and Supp Figure 6 for detailed cluster description).

Clusters M_6 and M_7 shared a similar expression profile characterized by hypoxic and angiogenic markers (*Vegfa*, *Hilpda*, *Bhlhe40*, *Hmox1*) together with a glycolytic gene expression pattern (*Slc2a1*), with M_6 showing higher expression of these genes. M_6 cells also expressed heat shock proteins (*Hspa1a*, *Hspa1b*), indicating their exposure to stress. M_7 was additionally expressing tissue remodelling genes, matrix metalloproteinases *Mmp12* and *Mmp13*, along with *Slc7a2* and *Il7r*. This gene profile is associated to cells previously described to predominantly exert an immunosuppressive role (44). Surprisingly, the frequency of M_7 subset appeared to be increased in the mice responding favourably to anti-PD-1 treatment (Figure 3C). Although there were no major changes in the overall proportion of subsets, closer examination revealed differentially expressed genes between progressors and regressors within specific subpopulations (Figure 3D and 3E). Subsets with the highest number of differentially expressed genes (DEGs) between the two groups were M_2 (monocytes), M_3 (Mono-macs), M_4 (MHC II^high^ TAMs) and M_5 (LA TAMs). The remaining subsets exhibited minimal or no differential expression of genes between the two groups. This considerable variability observed in the indicated subpopulations may indicate that cells from the two groups diverge the most in their functional states along the differentiation trajectory; not at their starting point (M_1) nor at their terminally differentiated states (M_6 and M_7). In three out of four analysed clusters, *Cd274* (encoding PD-L1), was consistently upregulated in regressors. Conversely, the gene *Mrc1*, which encodes the CD206 protein associated with suppression of anti-tumor immune responses, appeared to be downregulated within these identical subpopulations (Figure 3E).

The highest *Cd274* expression was observed in the M_7 cluster; however, a predominantly differential expression of *Cd274* emerged from M_3 mono-macs, which displayed increased PD-L1 expression in regressors (Figure 4A). In addition to upregulation of *Cd274*, M_3 subset in regressors was characterized by an increased expression of *Stat1*, *Tap1*, *Nampt* as well as *Gbp2* and *Gbp4*, known to be IFNγ-induced genes (45,46) (Figure 3E). Multiparametric flow cytometry approach confirmed the presence of a well-defined cluster of cells corresponding to previously identified mono-macs, characterized as F4/80^neg^Ly6C^pos^MHCII^pos^PD-L1^high^ cells (Figure 4B). In mice that responded to immunotherapy, there was a marked increase in the frequency of F4/80^neg^Ly6C^pos^MHCII^pos^ cells (Figure 4C, see Figure S2 for gating strategy). Furthermore, we confirmed an increased expression of PD-L1 in this cellular subset from regressor mice (Figure 4D). In a simplified analysis using a four-cluster model to compare different mono-macro subsets, both F4/80^neg^Ly6C^pos^MHCII^pos^ and F4/80^neg^Ly6C^low^MHCII^pos^ cells were characterized by elevated expression of CD40, CD80, PD-1, and PD-L1 (Figure S7), suggesting an enhanced antigen-presenting capacity linked to MHC II expression. In this simplified FACS approach, cells that were Ly6C^pos^MHC II^neg^ were identified as monocytes, corresponding to M_1/M_2 single-cell clusters. Ly6C^pos^MHC II^pos^ cells were considered to be transitionary monocytes, akin to the M_3 cluster, whereas Ly6C^low^MHC II^pos^ cells matched the MHC II^high^ TAM cluster identified by single-cell sequencing (cluster M_4). Cells that were Ly6C^neg^MHC II^neg^ were presumed to be a combination of various single-cell TAM clusters (M_5-M_7), generally associated with immunosuppressive properties.

**Figure 4.**
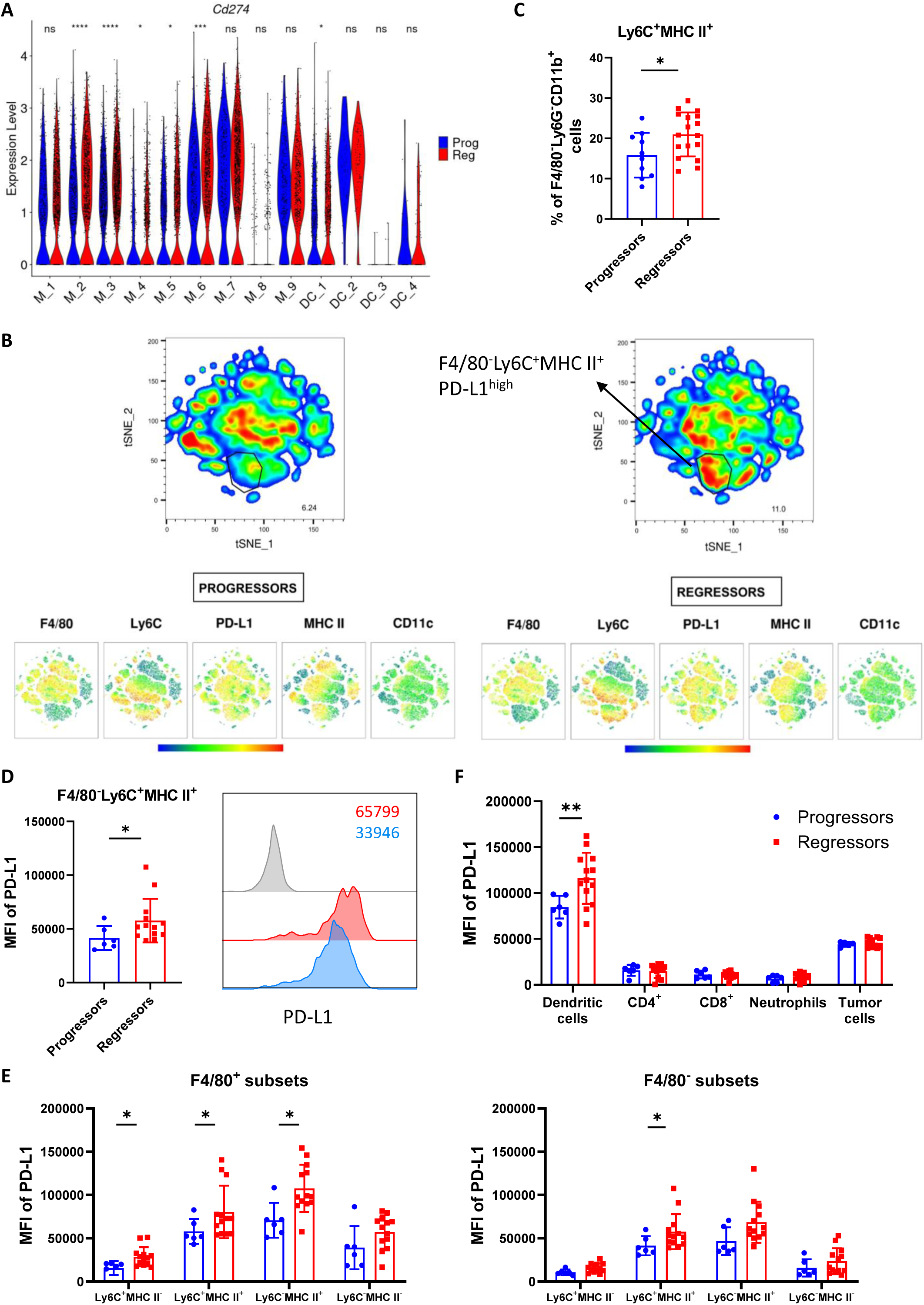
Pretreatment tumors of mice responding to immunotherapy display elevated expression of myeloid PD-L1. DBA/2 mice were inoculated s.c. with 1 x 10^6^ P815 mastocytoma cells in both flanks. Tumor infiltrating immune cells from the left flank were harvested 10 days after tumor inoculation and analyzed by flow cytometry or scRNA-seq. Mice bearing contralateral tumors were treated with 250 µg of anti-PD-1 monoclonal antibodies from day 10 to day 13 and tumor growth was followed. **A)** Violin plot of *Cd274* expression in identified single-cell clusters in progressors and regressors. **B)** t-distributed stochastic neighbor embedding (t-SNE) plot after dimensionality reduction and unsupervised clustering of flow cytometry data from CD11b^+^Ly6G^-^ myeloid cells from progressors (n=5) and regressors (n=5). **C)** Proportion of Ly6C^+^MHC II^+^ cells among F4/80^-^CD11b^+^Ly6G^-^ myeloid population in progressors and regressors. **D)** Quantification of MFI of PD-L1 in indicated subset and representative FACS histogram. **E)** MFI of PD-L1 in F4/80^+^ and F4/80^-^ myeloid subsets. **F)** MFI of PD-L1 in dendritic cells, CD4^+^, CD8^+^ T lymphocytes, neutrophils and tumor cells of progressors and regressors. Data show a pool of 3 or 4 independent experiments with 2-6 mice per group. Statistical significance was assessed by unpaired t test (D) and Mann-Whitney test (C, E). *p < 0.05, **p < 0.01. The absence of asterisks in the graph indicates no significant difference between groups.

Macrophages and DCs from responding mice were found to express higher levels of PD-L1 when compared to the progressor group. Notably, expression of this marker on T cells, neutrophils and tumor cells did not differ between mice responding or not to anti-PD-1 administration (Figure 4E and 4F). Collectively, these observations indicate that acquisition of PD-L1 by infiltrating monocytes at an intermediate stage of differentiation (MHC II^high^ clusters) could determine a positive response to PD-1-blockade immunotherapy.

### Pseudotime analysis reveals distinct differentiation dynamics of tumor-infiltrating myeloid cells from mice responding or not to anti-PD-1 treatment

To gain more insight into the potential differentiation pathways of the cell subpopulations identified, we performed trajectory analysis using the Slingshot R package (47). Trajectory inference was performed only on Mo-Mac cells to avoid any potential confounding factor. Five distinct trajectories were identified, all originating from M_1 cluster (monocytes that recently migrated from the bloodstream). Lineages 4 and 5 ended in clusters M_8 and M_9, respectively. However, lineages 1, 2 and 3 might be of potential significance due to the multiple differentiation states inferred along their trajectories with an early branching occurring within monocytic M_2 cluster (Figure 5A). Importantly, in the lineage 1 differentiation trajectory, TIMs (cluster M_1) first acquired features of transitionary monocytes (cluster M_3) before ultimately differentiating into the complete TAM subset that still retained MHC class II expression (cluster M_4). Cells following lineage 2 progressed from clusters M_1 and M_2 through the state of hypoxic TAMs (cluster M_6), and finally ended in a more differentiated TAM cluster M_7. Meanwhile, in the lineage 3, monocytic cells differentiated towards the *Mrc1*^high^ TAM cluster (cluster M_5), known for its immunosuppressive characteristics, with the minor fraction reaching the M_7 cluster. These findings reveal that upon infiltrating tumors, the majority of monocytes follow two main differentiation paths: one in which they acquire and maintain MHC class II expression; and another one where cells already diverge at the later monocyte stage, do not acquire MHC class II, but instead progress directly through the states with low MHC II expression. (M_1 → M_2 →M_5 or M_1 → M_2 → M_6 → M_7).

**Figure 5.**
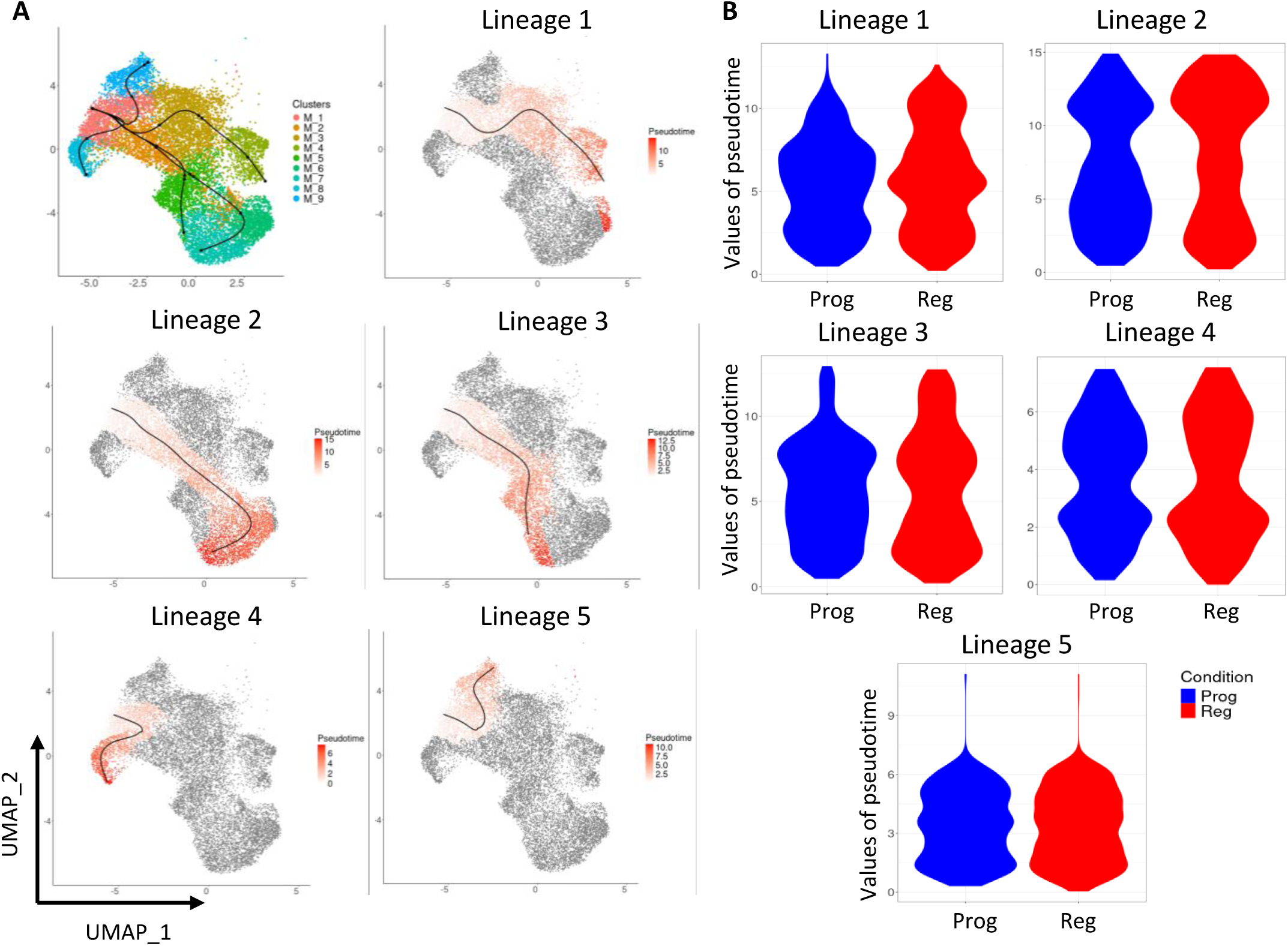
Tumor infiltrating myeloid cells from progressor and regressor mice undergo distinct differentiation dynamics. **A)** Trajectory analysis using Slingshot. Five lineages are shown together and separately on UMAP. **B)** Pseudotime analysis of five identified lineages between progressors and regressors.

To understand if there are any differences in cell progression along the identified trajectories between progressor and regressor mice, we compared the pseudotime values between these two groups. Lineage 1 differentiation trajectory showed a distinct profile between the two groups. In particular, most of the cells from mice unresponsive to immunotherapy displayed a deficit in cells reaching the final stage of the trajectory, suggesting that in progressor mice, monocytes failed to adequately acquire a more “mature” macrophage-like status. This observation also holds true for lineages 2 and 3, where cells from regressors appeared to reach more advanced differentiation states. Lineages 4 and 5 displayed no major differences along their differentiation trajectories between progressors and regressors (Figure 5B). Collectively, differences in distribution of pseudotime values between the two groups suggest that TIMs from progressor and regressor groups undergo distinct differentiation dynamics, likely influenced by environmental factors. Cells in progressors may be arrested at an earlier stage of differentiation or this could also reflect a variation in the timing of differentiation among the cells from the two groups.

### IFNγ-dependent expression of PD-L1 by myeloid cells in mice responding to immunotherapy

To further understand which environmental factor may be responsible for the elevated expression of PD-L1 in specific cell subsets and distinct differentiation dynamics observed between the two groups, we performed upstream regulator and pathway analysis on differentially expressed genes. This functional analysis highlighted pathways linked to cytokine signaling and various immune responses as playing an important role in defining animals responding to immunotherapy (Figures 6A and 6B). Collectively these data suggest that rather than the overrepresentation of a particular cell subset, the TME of regressor mice was characterized by the response to a cytokine-enriched milieu. IFNγ has been previously shown to positively regulate expression of PD-L1 and PD-L2 (48). To evaluate the role of IFNγ on tumor growth and expression of PD-L1, mice were inoculated with 2x10^6^ P815 cells in a single flank and treated with blocking anti-IFNγ (Figure 6C). As expected, in vivo blocking of this cytokine led to increased tumor growth (Figure 6D). Administration of anti-IFNγ mAbs led to a selective decrease of PD-L1 expression by infiltrating immune cells, while leaving tumor-associated expression of this checkpoint protein unaffected (Figure 6E). The effect of IFNγ blocking was more pronounced in CD11b-expressing cells, where PD-L1 expression levels were substantially reduced, whereas T cells appeared insensitive to this treatment (Figure 6E and 6F), further confirming the predominant influence of PD-L1 expression by myeloid cells. Of note, in addition to the diminished PD-L1 expression, inhibiting IFN-γ also resulted in a decreased expression of other proteins typically induced by IFN-γ, such as Sca-1 (encoded by the gene *Ly6a*, Figure S8A). Importantly, blocking IFNγ led to a selective and significant increase in the frequency of CD11b^pos^F4/80^pos^ cells (and in particular the Ly6C^neg^MHCII^neg^ subset, generally associated to TAMs, see Figure 6G and S8B). In contrast, the same treatment caused a significant loss of cells co-expressing Ly6C and MHC II, as well as MHC II^high^ cells (Figure 6H and Figure S8B). Although the anti-IFNγ treatment did not significantly alter the phenotype of CD8^+^ TILs (Figure S8E), it led to a slight increase in CD8^+^ tumor infiltrating cells, with a preferential expansion of cells reactive to the P1E antigen (Figure S8D). A spontaneous response characterized by the production of IFNγ appears therefore to be responsible for the accumulation of myeloid cells (including DCs, see Figure S8C) whose frequency and phenotype were found to be associated to a regressor status.

**Figure 6.**
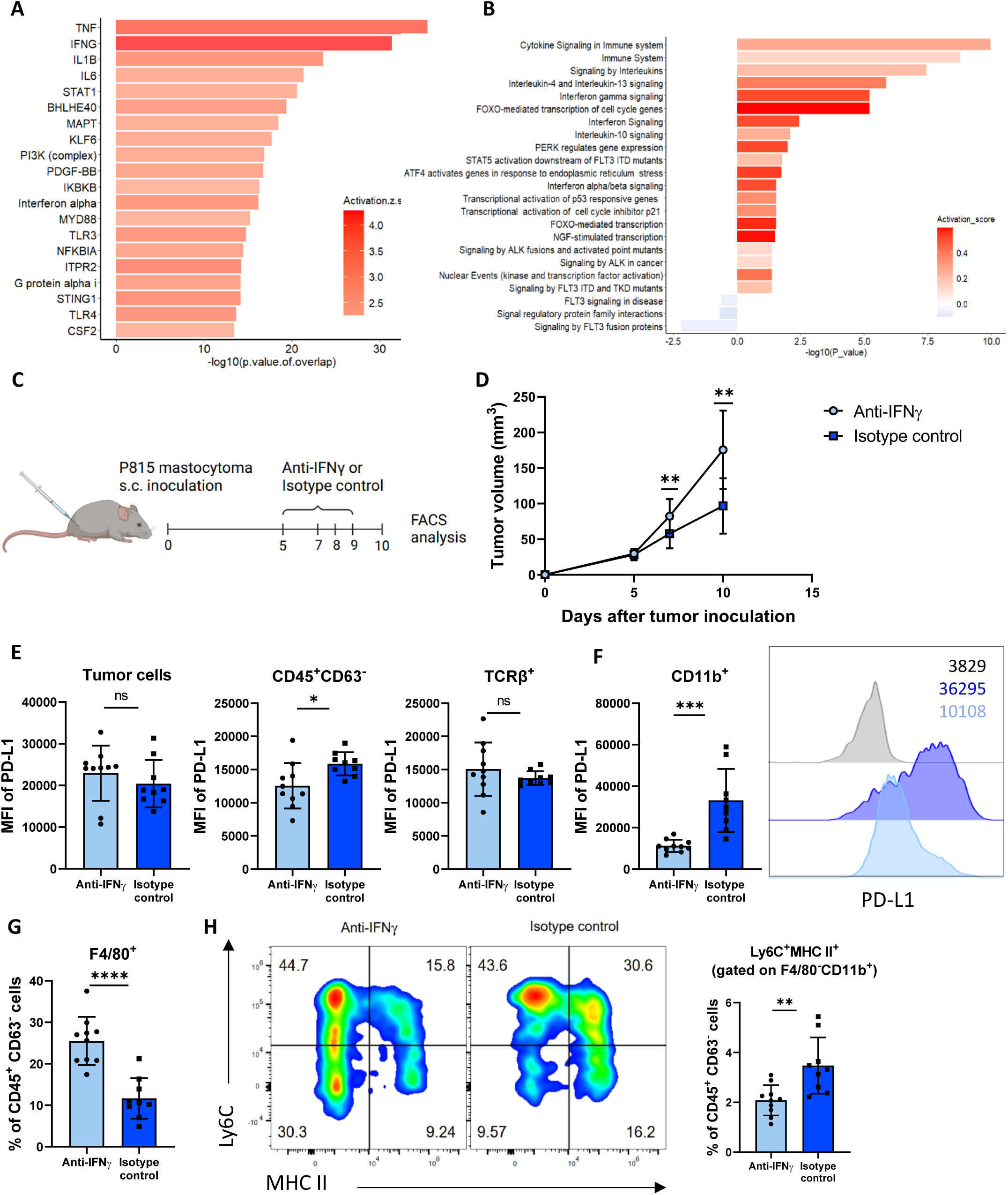
PD-L1 expression in myeloid cells is IFNγ-dependent. **A)** Upstream regulators of genes upregulated in regressors predicted by Ingenuity Pathway Analysis (IPA). **B)** Gene set enrichment analysis (GSEA) on DEGs in progressors versus regressors using Reactome Pathway Database. **C)** Schematic of the experimental set-up. DBA/2 mice were inoculated s.c. with 2 x 10^6^ P815 cells in a single flank and treated intraperitonealy with 250 µg of anti-IFNγ monoclonal antibodies or isotype control. Tumors were harvested 10 days after tumor inoculation and analyzed by flow cytometry. Created with BioRender.com. **D)** Tumor growth of P815 mastocytoma. MFI of PD-L1 in **E)** tumor cells, CD45^+^ immune cells, TCRβ^+^ population and **F)** CD11b^+^ myeloid cells with representative FACS histogram. **G)** Proportion of F4/80^+^ TAMs among CD45^+^CD63^-^ immune population. **H)** Representative FACS plot and proportion of Ly6C^+^MHC II^+^ subset among CD45^+^CD63^-^ immune cells. (D-H) Data are representative of 1 experiment with 9-10 mice per group. Statistical significance was assessed by Mann-Whitney test or unpaired t test. *p < 0.05, **p < 0.01, ***p < 0.001, ****p < 0.0001.

## DISCUSSION

Although the presence of immune cells within the tumor bed appears to be a prerequisite for a favourable response to ICB, not all patients presenting with an inflamed tumor respond to these treatments. The aim of the present study was to develop and exploit a murine model of ICB to identify the nature and quality of immune cell infiltrates that can be associated with a positive anti-tumor response following the administration of anti-PD-1 monoclonal antibodies.

This study was based on the observation that DBA/2 mice implanted with P815 mastocytoma displayed a dual response to PD-1 blockade, in which only a proportion of the animals (on average 50%) rejected their tumors. This dual response was also observed in mice in which the cancer cell line was implanted subcutaneously into both flanks, providing an opportunity to explore the influence of the pretreatment TME on the response to ICB. Using a combination of single cell-based techniques (flow cytometry and single-cell RNA-seq), we were able to identify correlates of tumor regression in response to anti-PD-1 treatment. When analysed globally, no statistically significant differences in the abundance of T, NK, or myeloid cell types were found between responders and non-responders. However, after sub-clustering for T cell and myeloid cell populations, distinct features in the TME of mice responding to ICB could be identified.

Not surprisingly, and in agreement with numerous animal and clinical studies, an increased frequency of CD8^+^ T cells was found to be associated with a positive response to ICB (49–51). Taking advantage of the previously identified P815 tumor-specific antigens, we were able to better define the prerequisite of CD8^+^ T cell infiltration that was best associated with a productive anti-tumor response. The presence of a minor population of lymphocytes specific for a tumor-associated self-antigen (P1A) was a better predictor of response when compared to the frequency of cells specific for a P815-neoantigen (P1E). When compared to the total CD8^+^ infiltrating population (presumably comprising antigen-non-reactive, passenger lymphocytes), P1A-reactive cells displayed a phenotype associated with TCR signalling and an exhausted state, such as increased expression of PD-1, CD38, Tim-3 (Fig 2G), and TOX (Fig S3B). When the phenotype of P1A tetramer^+^ cells was compared between progressor and regressor mice, Tim-3 and in a less marked fashion, CD38 expression appeared to be a major phenotypic marker associated with tumor regression in response to anti-PD1-based therapy. In agreement with this conclusion, despite their lower frequency, Tim-3 expression was also found to be elevated among P1E-reactive cells in responding mice (Fig 2G). Based on this finding, we analysed the functional properties of CD8^+^Tim-3^+^ cells, irrespective of their antigen specificity, to identify potential correlates of response. This analysis revealed that increased cytotoxicity (based on granzyme B and cell surface CD107a expression) and IFNγ-secreting potential were associated with high levels of Tim-3 expression. This apparent contrast between an exhausted-like phenotype and effector function potential is not without precedent. Single-cell analyses of CD8^+^ T lymphocytes infiltrating human lung cancer masses revealed the presence of PD-1^high^Tim-3^high^CD38^pos^ cells expressing high levels of cytokines and cytotoxic factors (52). The presence of CD8^+^ T cells displaying a tissue-resident phenotype (including PD-1 and Tim-3 expression) was found to be associated with a positive response to anti-PD-1 therapy among lung cancer patients (53) and metastatic melanoma patients (54). Finally, a recent study performed on a murine model of haematopoietic malignancies (55) revealed the presence of CD8^+^ T cells that despite expressing an exhausted phenotype, retained full functional potential as judged by cytotoxic and IFNγ secretion capacities. Taken collectively, these studies, including our own, call for caution in establishing a strong causal correlation between the expression of negative costimulatory molecules and a lack of response to ICB.

Although additional efforts would be required to better understand the predominant role played by a minor, self-reactive T lymphocyte population in determining a positive response to ICB, recent studies aimed at exploring the role of TCR avidity on a successful response to immunotherapy may provide a plausible explanation. Several publications concur with the idea that lymphocytes expressing TCR of lower antigen-binding strength/avidity provide better protection against chronic antigen exposure. High-signal-strength interactions result in T cell dysfunction and exhaustion, whereas low-signal-strength interactions cause a less marked form of T cell functional inertness (56). Similarly, CAR-T cells expressing high affinity TCRs may be more prone to terminal exhaustion (57), while lower affinity CAR T cells display increased proliferation and in vivo antitumor activity (58,59). Accordingly, PD-1 appears to preferentially inhibit the activation of low-affinity T cells (60), which is in line with several observations indicating that the immunotherapeutic efficacy of anti-PD-1-based ICB is associated with the reactivation of low avidity, tumor-specific lymphocytes (61,62). Although we cannot evaluate the overall avidity of P1A vs P1E-specific lymphocytes in our model (these tumor antigens are presented by distinct MHC molecules whose tumor-cell surface MHC-peptide complex density remains to be established), it is noteworthy that neoantigen specific T cells (P1E-reactive) display a reduced TCR expression in the draining lymph nodes (a privileged site of anti-tumor response generation, (63)), compatible with a sustained, and possibly high avidity interaction of these cells with their cognate Ag/MHC complex.

We next analysed in more details the tumor-infiltrating myeloid cell compartment, comparing the frequency of several subpopulations of innate cells between progressor and regressor mice. As expected, monocyte and monocyte-derived cells represented a sizable population of TME-associated immune cells. The functional properties and density of these cells within the tumor bed has been clearly shown to play a role in the response to ICB (64). The advent of single-cell based technologies has revealed the extensive plasticity and therefore functional heterogeneity of these monocyte-derived cells, well beyond the classical M1 vs M2 dichotomy. It is presently too early to firmly attribute a clear cell identity to all the tumor associated monocyte/macrophage subpopulations, and the field awaits further analyses to achieve a consensual terminology and functional characterisation of these cells. However, trajectory inference and pseudotime analyses offer the possibility to align cells along a putative differentiation pathway, based on gradual differences in their transcriptome. Rather than attempting a full characterisation of TAMs in our model, we have focused our analyses on cell phenotype and trajectories that differ between mice responding or not to ICB. Single-cell RNAseq of tumor infiltrating myeloid cells revealed the presence of 9 subpopulations of TAMs and 4 DC subsets in the TME. Because of their large representation, we mainly focused our attention on monocytes and monocyte derived cells.

We identified three trajectories of interest that exhibit the most prominent differences between progressors and regressors, suggesting two possible differentiation paths upon monocyte infiltration into the TME. The first lineage we considered leads early monocyte (M_1 subset, displaying very few DEGs among progressor and regressor mice, in agreement with their recent developmental history) to a subset of cells (M_4) characterized by high expression of complement-associated factors and MHC II molecules. Pseudotime analysis suggests an increased propensity of cells from regressor mice to reach this MHC II^high^ status (see Figure 5B). The second major path (represented by lineages 2 and 3) leads to a population of cells expressing a “regulatory” phenotype, characterized by the expression of *Arg1*, *Hmox1, Cd274* (PD-L1) associated to a low, but detectable MHC II expression (see Fig S6). Frequency of these “regulatory TAMs” appears to be elevated in regressor mice (Figure 3C). Thus, rather than being restricted to a single cell population, this analysis revealed a possible dual differentiation path (mostly lineage 1 and 2-3) that differ from regressor and progressor mice. PD-L1 expression was consistently upregulated in clusters along lineage 1 (the lineage of MHC II^high^ states) in regressor mice. This observation suggests that PD-L1 expression, together with MHC II, represents an important marker for predicting responsiveness to anti-PD-1 therapy. Although caution must be exerted when comparing transcriptomic data to protein-based assays, flow cytometry analyses confirmed the persistence of MHC II positive cells and cells expressing PD-L1 (often expressed by the same cells) as a hallmark of mice responding to ICB. Taking advantage of the transcriptome analysis performed on the unseparated myeloid subset, we identified IFNγ as a possible upstream regulator whose influence is more pronounced in regressor mice. This cytokine is responsible for the elevated expression of PD-L1 and MHC II molecules on subsets of myeloid cells, as illustrated by the analyses of tumor infiltrating cells isolated from mice injected with blocking anti-IFNγ antibodies (Figure 6). Notably, this experiment revealed an important role for this cytokine in supporting PD-L1 expression by myeloid, but not by tumor or tumor-infiltrating lymphocytes. Although merely correlative, this observation concurs with recent finding highlighting the important, and often predominant, role of PD-L1 expressed by myeloid vs tumor cells in determining a therapeutic response to ICB (65–69). These observations are also in line with clinical findings indicating that an IFNγ-related gene signature was associated to a therapeutic response to anti-PD-1-based ICB (70–72).

Collectively, these single cell analyses suggest that responsiveness to ICB may also be impacted by the differentiation dynamics of monocyte-derived TAMs at the onset of the treatment. This emphasizes the necessity to decipher the epigenetic landscape underlying these differences and direct future research towards a temporal analysis of TAMs. Identifying the factors responsible for transition between differentiation states in the TME could enable us to further advance the cells along their predetermined differentiation trajectory, thereby potentially enhancing the response to ICB (73–75).

Despite continuous growth, all tumors analysed in this study can be considered as “hot” or “inflamed” based on the presence of infiltrating immune cells. We previously demonstrated that T lymphocytes in this model can limit, but not fully impede tumor growth (76). A spontaneous, yet inefficient, immune response characterized by an IFNγ signature only develops in a subset of mice, possibly in a stochastic fashion. Based on our analyses, IFNγ appears as mainly produced by infiltrating, tumor-specific CD8^+^ T cells. Although all infiltrating CD8^+^ T lymphocytes may contribute to the secretion of this pro-inflammatory cytokine, our study also reveals that not all tumor reactive T cells may actively contribute to tumor rejection after administration of anti-PD-1 mAbs. It is tempting to speculate that moderate TCR-tumor Ag interactions may lead to a reversible state of exhaustion, representing a privileged source of effector cells upon release from PD-L1-dependent negative signalling. Secretion of IFNγ will likely promote/sustain expression of MHC II and PD-L1 by a subset of TAMs, which in turn may both support (via MHC II presentation to CD4 cells) and control (through PD-L1 expression) an anti-tumor response. Although often correlative, our observations and the existing literature suggest therefore that a positive response to anti-PD-1 ICB may be linked to a positive feedback loop involving IFNγ producing TILs and MHC II^pos^PD-L1^pos^ monocyte derived macrophages (77).

We acknowledge several limitations to our study. Despite numerous attempts and the existing literature (78), we could not identify a mouse tumor model in the B6 genetic background displaying a similar dual response to ICB, limiting therefore a genetic validation of some of our findings. The role of TCR avidity and/or density and nature of tumor antigens in regulating T cells responsiveness to anti-PD-1-based ICB remains unexplored. A more comprehensive phenotyping (i.e. by single cell RNAseq) of P1A vs P1E - reactive T cells may contribute to a better understanding of the signalling pathways that sustain T cell effector function recovery after ICB. Moreover, the relative contribution of tumor-infiltrating vs lymph node-associated T lymphocytes has not been evaluated in this study. The consequences of IFNγ blocking on the differentiation path of monocyte-derived cells should be explored in a more comprehensive fashion, using single cell RNAseq and spatial transcriptomics.

In any event, studies like ours and others (35–37), based on reductionist mouse models approach, suggest that response to ICB can vary independently of tumor and host genetic factors. In addition to provide valuable markers to identify patients more likely to respond to immunotherapy, these approaches may suggest combinatory strategies aiming at augmenting in situ IFNγ production to increase patient sensitivity to ICB.

## MATERIAL AND METHODS

### Mice

DBA/2 Ola Hsd mice were purchased from Envigo (Horst, The Netherlands) and bred in the animal facility of Biopark ULB Charleroi, Belgium (BUC). Mice were bred and housed under specific pathogen-free conditions (SPF). Mice used for the experiments were females between 7 and 12 weeks of age. Experiments were carried out in accordance with the relevant laws and institutional guidelines and approved by the ULB Institutional Animal Care and User Committee.

### Cell culture

Murine mastocytoma cell line P815 (clone P1.HTR-3), and the P1A KO cell line (P1.204), a variant of P815 carrying a deletion of gene *Trap1a*, have been previously described (79,80). Tumor cells were cultured in DMEM (Lonza) supplemented with 10% fetal bovine serum (FBS, Sigma-Aldrich), 1% sodium pyruvate, 1% L-glutamine, 1% non-essential amino acids (all reagents from Lonza) and 0.1% β-mercaptoethanol (Sigma-Aldrich) at 37 °C and 5% CO2. Cell medium was changed every 2 days. Cells were regularly checked for the presence of mycoplasma contamination.

### In vivo experiments

Cells were harvested in the exponential growth phase and resuspended in sterile PBS for inoculation. DBA/2 mice were injected subcutaneously (s.c.) in one flank with 2 x 10^6^ P815 mastocytoma cells or bilaterally (in both flanks) with either 1 x 10^6^ P815 cells or P1A KO cells. Tumor size was measured three times per week and tumor volume was determined as (length x width^2^)/2. Only mice harbouring well-established tumors, as determined by a tumor volume falling within the range of 100 to 200 mm^3^, and with similarly sized tumors on each flank, were enrolled in the experiment. Mice were monitored for at least 50 days after tumor inoculation and euthanized when humane endpoint was reached.

Mice were treated intraperitoneally (i.p) with anti-PD-1 anti-mouse monoclonal antibody (clone RMP1-14, either purchased from BioXcell, USA, or produced internally, with similar outcomes) at 250 µg/mouse every day from day 10 to 13 or with isotype control (anti-β-Gal, clone GL117). Treatment was initiated the same day as surgery. Mice were injected i.p. with anti-IFNγ monoclonal antibody (clone XMG1.2, BioXcell) or IgG1 isotype control (clone HRPN, BioXcell) at 250 µg/mouse at day 5, 7, 8 and 9 after tumor inoculation.

### Surgical operation

Mice were anesthetized by i.p. injection of a combination of 5% xylazine and 10% ketamine prepared in sterile PBS (10 µl/g). Tumor was removed through an incision in the flank and immediately immersed in cold PBS. Surgical wound was closed with suture clips. Mice were designated as regressors when their tumors completely regressed and they remained tumor free for at least 5 weeks after treatment. Mice were designated as progressors when their tumor growth was not impacted by the treatment, as in isotype control group.

### Cell preparation

For all experiments, organs were processed under sterile conditions. Tumors were perfused with a solution containing DNase I (Roche) and Liberase (Roche), cut in smaller pieces and incubated for 30 min at 37°C. The suspension was washed with RPMI (Lonza), 5% FBS and 2 mM ethylenediaminetetraacetic acid (EDTA, VWR), mashed and filtered through a 70 μm cell strainer, twice. Lymph nodes were mashed, filtered and washed in HBSS (Lonza). Spleen were mashed and red blood cells were lysed by ACK lysing buffer (Ammonium-Chloride-Potassium, home-made). Suspension was neutralized by adding RPMI, 5% FBS and filtered before further manipulation.

### Flow cytometry

Single-cell suspensions were stained with live/ dead fixable near-IR dead cell stain kit (Thermo Fisher Scientific) or iFluor 860 maleimide (AAT Bioquest) to exclude dead cells, incubated with Fc receptor-blocking antibodies CD16/32 (clone 2.4G2, BioXcell) to block non-specific binding and stained with the following fluorochrome-conjugated antibodies for surface markers: CD45 (30-F11), CD45.2 (104), TCRβ (H57-597), CD8α (53-6.7), CD4 (GK1.5 or RM4-5), CD11b (M1/70), CD11c (HL3), F4/80 (T45-2342), MHC II (I-A/I-E, M5/114.15.2), CD19 (1D3), Ly6C (AL-21), Ly6G (1A8), PD-L1 (MIH5), CD44 (IM7), CD69 (H1.2F3), CD38 (90/CD38), Tigit (1G9), Tim-3 (5D12/TIM-3), PD-1 (J43), CD62L (MEL-14), CD3 (17A2), NKp46 (29A1.4), CD49b (DX5) from BD Biosciences, CD63 (NVG-2), CD80 (16-10A1), CD107a (1D4B), CD40 (1C10) from Thermo Fisher Scientific and Sca-1 (D7) from BioLegend. For tetramer staining, cells were stained with P1A/H-2L^d^-PE or P1E/H-2K^d^-PE tetramer. Tetramer complexes were produced as previously described (81). For detecting intranuclear proteins, cells were fixed and permeabilized for 30 min with eBioscience Foxp3/Transcription Factor Staining Buffer Set (Thermo Fisher Scientific) before staining with the following antibodies: Eomes (Dan11mag), T-bet (4B10) and TOX (TXRX10) from Thermo Fisher Scientific.

To evaluate the functional status of tumor-infiltrating lymphocytes, cells were stimulated for 4h with 50 ng/ml of phorbol 12-myristate 13-acetate (PMA, Sigma-Aldrich) and 1 µg/ml of ionomycin calcium salt (Sigma-Aldrich) in the presence of brefeldin A (Thermo Fisher Scientific). Afterwards, cells were stained for surface markers (see above), followed by fixation with BD Cytofix/Cytoperm and permeabilization with BD Perm/Wash (BD Biosciences). Cells were stained with conjugated monoclonal antibodies directed at IFNγ (XMG1.2) from BD Biosciences, Granzyme B (NGZB) and TNFα (MP6-XT22) from Thermo Fisher Scientific or Granzyme B (GB11) from BD Biosciences.

Flow cytometry data were acquired on a CytoFLEX LX (Beckman Coulter Life Sciences) and analyzed using FlowJo software (v10.8.1).

### Single-cell RNA sequencing

#### Library preparation, sequencing and alignment

Single-cell suspensions from tumors were obtained as described. Cells were labelled with TotalSeq-B anti-mouse hashtag antibodies and at the same time stained with an antibody mix used for sorting cell population of interest. Five different TotalSeq-B antibodies were used in each experiment (Hashtag antibody 1-5, BioLegend). Incubation lasted for 30 min. LIVE/DEAD^-^CD63^-^TCRβ^-^CD19^-^ CD45.2^+^Ly6G^-^CD11b^+^CD11c^+^ cells were sorted from tumor cell suspensions using a BD FACSAria III. After sorting, cell suspensions were filtered and pooled before being centrifuged for 10 min at 300g. Pellets were resuspended in RPMI 10% FBS medium and loaded on the Chromium Controller (10x Genomics). In total, cells from 6 progressors and 9 regressors were sequenced. Single-cell RNA-seq libraries were prepared using the Chromium Single Cell 3’ v3.1 Reagent Kit (10x Genomics) according to manufacturer’s user guide CG000206. Libraries were loaded to an Illumina Novaseq (Brightcore platform) and Cell Ranger (10x Genomics) functions mkfastq and count were used to demultiplex the sequencing data, generate gene-barcode matrices, and align the sequences to the mm10 genome. Resulted matrices were used to generate a single seurat object using Seurat 4.2.0 R package.

#### Demultiplexing

Data containing the information about oligo-tagged antibodies and cell barcodes were added as an independent assay in the Seurat object and were normalized with the NormalizeData function from the Seurat package (version 5.0.3), using the “CLR” normalization method which applies a centered log ratio transformation. The HTODemux function was used to reassign individual cells to their original mouse sample. Negative cells, with no oligo-tagged antibody detected, were removed.

#### Filtering

RNA data were filtered by mouse sample with the following thresholds: number of genes expressed > 1000 (between 800 and 1200 depending on the mouse sample), fraction of mitochondrial content < 20%, and total counts per cell > 1800 (between 1600 and 2400). The new single-cell gene expression profiles were normalized with the NormalizeData function using the “LogNormalize” normalization method and were scaled with the ScaleData function from Seurat package. For each biological replicate, the top 2000 variable genes in each condition were identified using the “vst” method implemented in Seurat. These variable genes were used to perform a principal components analysis (PCA) on all samples. The top 20 PCA components were selected to integrate the data with harmony (version 1.2.0) with sample batch as confounding factors (max.iter.cluster = 20) (82). Integrated components were then used to build a Uniform Manifold Approximation and Projection (UMAP) embedding of the 14,585 integrated cells (4,575 cells from progressor samples and 10,010 cells from regressor samples). Unsupervised clustering of the integrated data was performed with the Louvain algorithm to determine the different cell types (resolution = 0.8, k = 10).

#### Heatmap generation

Five marker genes upregulated in each population with the highest fold change were represented as a heatmap using the DoHeatmap function from the Seurat package. Distribution of cell percentages among regressors and progressors in a boxplot was constructed using the ggplot2 package (version 3.5.0).

#### Trajectory analysis

The Slingshot package (version 2.6.0) was used to perform trajectory analysis on the integrated Seurat object without dendritic cells in order to remove confounding factors. Lineages were calculated after selecting the “M_1” group as the root, as these cells express classical monocyte genes.

#### DEG analysis

Genes differentially expressed between regressors and progressors were determined using the FindMarkers function of the Seurat package and the MAST method from the MAST package (version 1.24.1). Genes with an adjusted p-value of 0.05 and with a logFoldChange absolute value greater than 0.25 were retained and represented as a volcano plot using the ggplot2 package. The same analysis was performed for the DEGs per cell population.

#### Upstream regulator and pathway analysis

Upstream regulator analysis was performed with IPA (Ingenuity Pathway Analysis from Qiagen) based on the list of DEGs. IPA calculates an activation score for key regulatory genes based on the fold change of the DEGs. Upstream regulators with an activation score greater than 2 and a p-value less than 0.05 were retained. Pathway analyses were performed with Reactome (83). Barplots were performed using the PPInfer package (1.20.4).

#### Statistical *test*

Statistical analysis of the cell percentages visualized in the boxplot was carried out using the stat_compare_means function from the ggpubr package (0.6.0) with the wilcox.test method performing a Wilcoxon-Mann-Whitney test.

### Statistical analysis

Data are presented as mean ± SD. Statistical analyses were performed using GraphPad Prism software (8.0.2). Difference between the experimental groups was assessed using two-tailed unpaired t-test or two-tailed unpaired Mann–Whitney test for comparing two groups. One-way ANOVA with Tukey’s multiple-comparison test or Kruskal–Wallis test with Dunn’s multiple-comparison test was used for comparing more than two groups. For evaluating the correlation between variables, two-tailed Spearman correlation analysis was used. Tumor rejection rate was assessed by two-sided Fisher’s exact test. A p value < 0.05 was considered significant. In figures, asterisks denote statistical significance as follows: *, p < 0.05; **, p < 0.01; ***, p < 0.001; ****, p < 0.0001.

## ACKNOWLEDGMENTS

We thank Valérie Acolty, Caroline Abdelaziz and Véronique Dissy for animal care and for technical support. We acknowledge the help of Frederick Libert and Anne Lefort for the sequencing experiments and bioinformatic analyses. This work was supported by the National Fund for Scientific Research (FNRS), the European Regional Development Fund (ERDF), the Walloon Region (ERDF-Wallonia BioMed 2014-2020-LIV 45-20), the Fondation contre le Cancer and by grants from the Fonds Jean Brachet and the Fondation Hoguet. J.G. and S.V.V have been supported by a Télevie grant from the FRS-FNRS, and S.G. is a research director of the FRS-FNRS.

## AUTHOR CONTRIBUTIONS

J.G. performed most of the experiments, analysed the data and prepared the figures. S.V.V. performed all bioinformatics analysis and prepared the relevant figures. C.H. helped in the study design, developed the original bilateral tumor model and performed the initial set of experiments. S.D. helped in the development of most flow cytometry studies and performed some experiments. A.A. performed key experiments related to the single cell studies. L.B. and B.J.V.D.E provided valuable reagents. M.M. designed and supervised the study. O.L. designed and supervised the study, provided funding and wrote the manuscript with the help of J.G. and S.G. S.G. supervised the study and provided funding.

---

**Figure S1.**
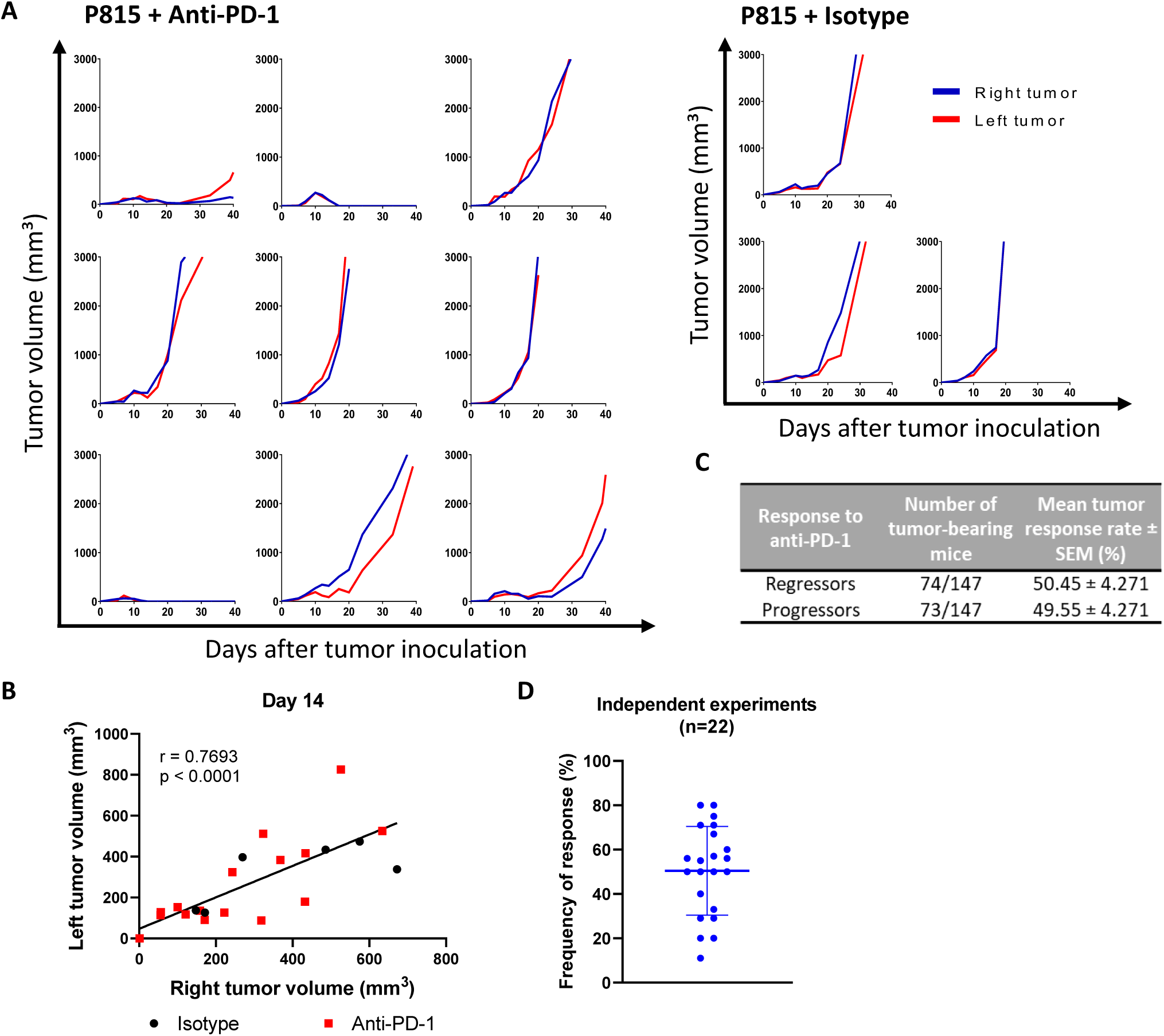
Response to anti-PD-1 treatment is a property of individual mice. DBA/2 mice were inoculated s.c. with 1 x 10^6^ P815 mastocytoma cells in each flank and 10 days later treated i.p. with 250 µg of anti-PD-1 or isotype control (from day 10 to day 13) respectively. Tumor volume was measured 3x per week. **A)** Growth of P815 tumors treated with anti-PD-1 or isotype control. **B)** Correlation of bilateral P815 mastocytoma tumor volumes 14 days after inoculation in anti-PD-1-treated or isotype-treated mice. **C)** Mean tumor response rate ± SEM and **D)** frequency of response to anti-PD-1 treatment. Each point represents an independent experiment. A) Data are representative of 1 experiment. B) Data show a pool of 2 independent experiment with 3-8 mice per group or (C-D) a pool of 22 independent experiments with 4-11 mice per experiment. Correlation analysis was performed using Spearman correlation (B).

**Figure S2.**
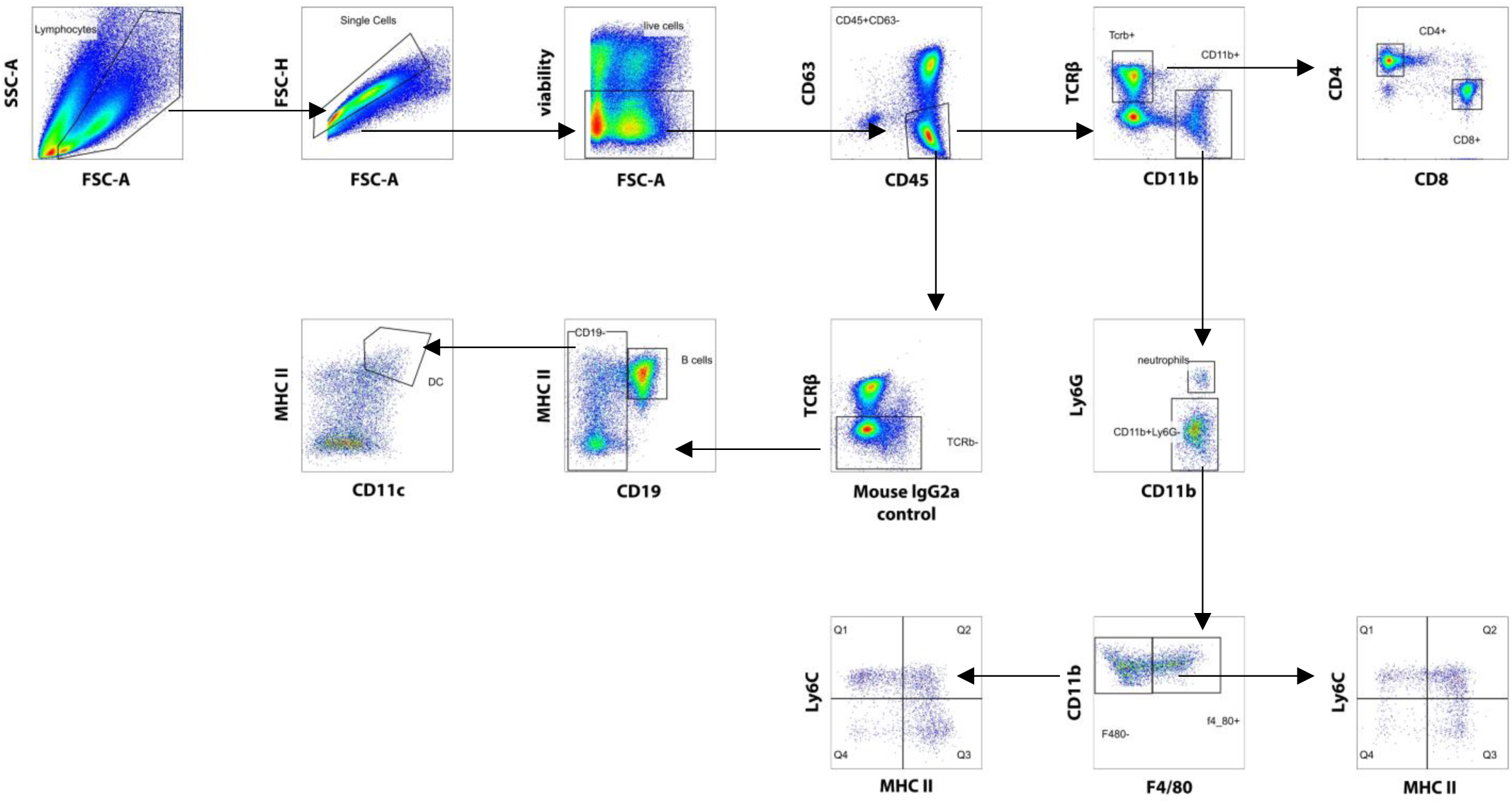
Flow cytometry gating strategy to analyze the influence of tumor-infiltrating immune cells on the response to immunotherapy.

**Figure S3.**
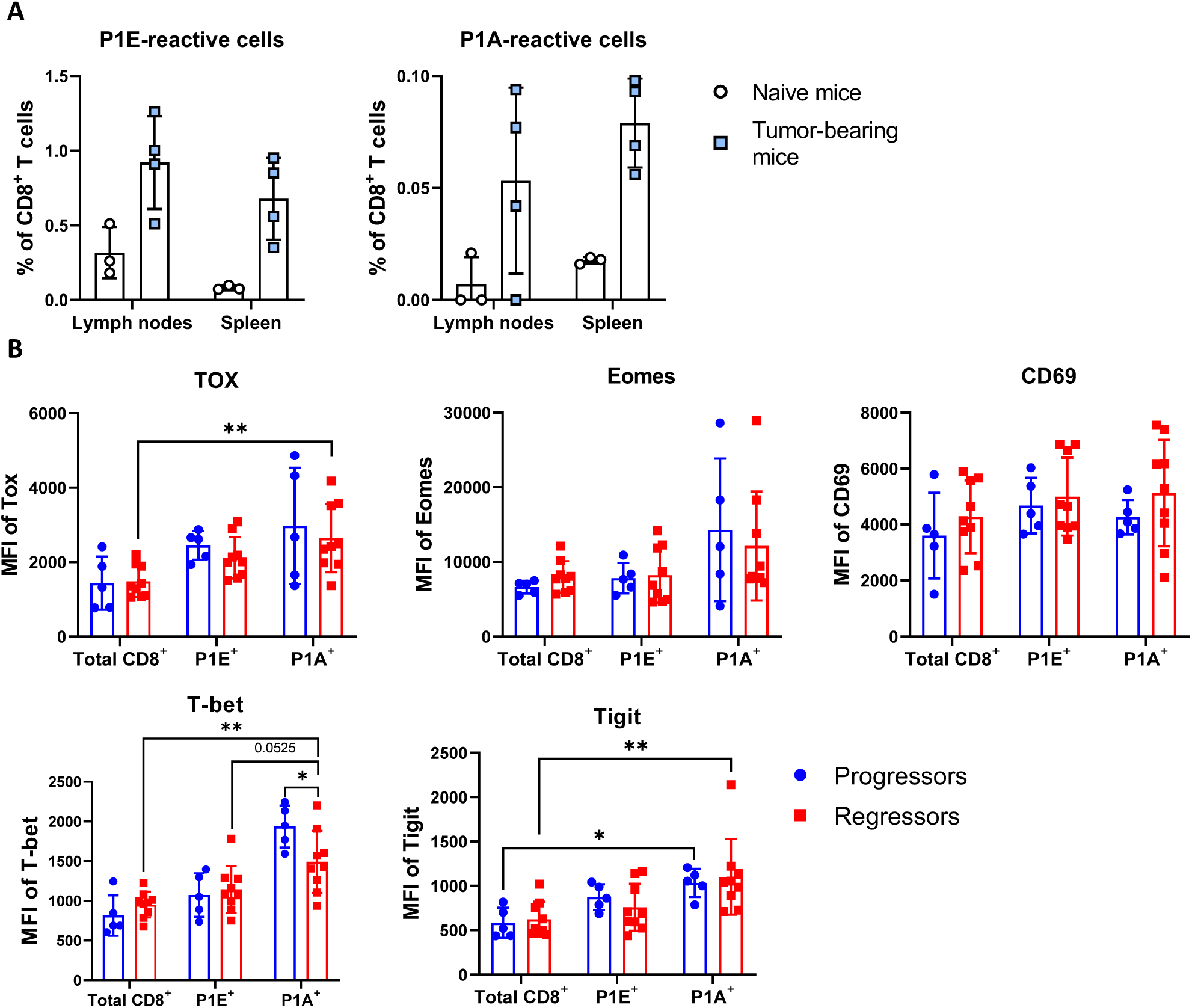
Phenotypic characterization of P1E- and P1A-specific CD8^+^ T cells. DBA/2 mice were inoculated s.c. with 1 x 10^6^ P815 mastocytoma cells in both flanks. Inguinal lymph nodes and spleen were collected 10 days later and frequency of tumor-reactive CD8^+^ T cells was examined by flow cytometry. **A)** Frequency of P1E- and P1A-reactive cells among CD8^+^ population in tumor-bearing and naive mice. Naive mice are animals that were not implanted with P815 mastocytoma cells. **B)** Tumors from one flank were surgically extracted 10 days after tumor inoculation and tumor-infiltrating immune cells analyzed by flow cytometry. Mice bearing contralateral tumors were treated with 250 µg of anti-PD-1 monoclonal antibodies from day 10 to day 13 and tumor growth was followed. MFI of TOX, T-bet, Eomes, Tigit and CD69 in CD44^+^ P1E- and P1A-specific CD8^+^ T cells in progressors and regressors. Data are representative of A) 1 experiment with 3-4 mice per group or B) show a pool of 2 independent experiments with 2-5 mice per group. Statistical comparisons were performed by using one-way ANOVA with Tukey’s multiple comparisons test or Kruskal-Wallis test with Dunn’s multiple comparisons test (B). *p < 0.05, **p < 0.01.

**Figure S4.**
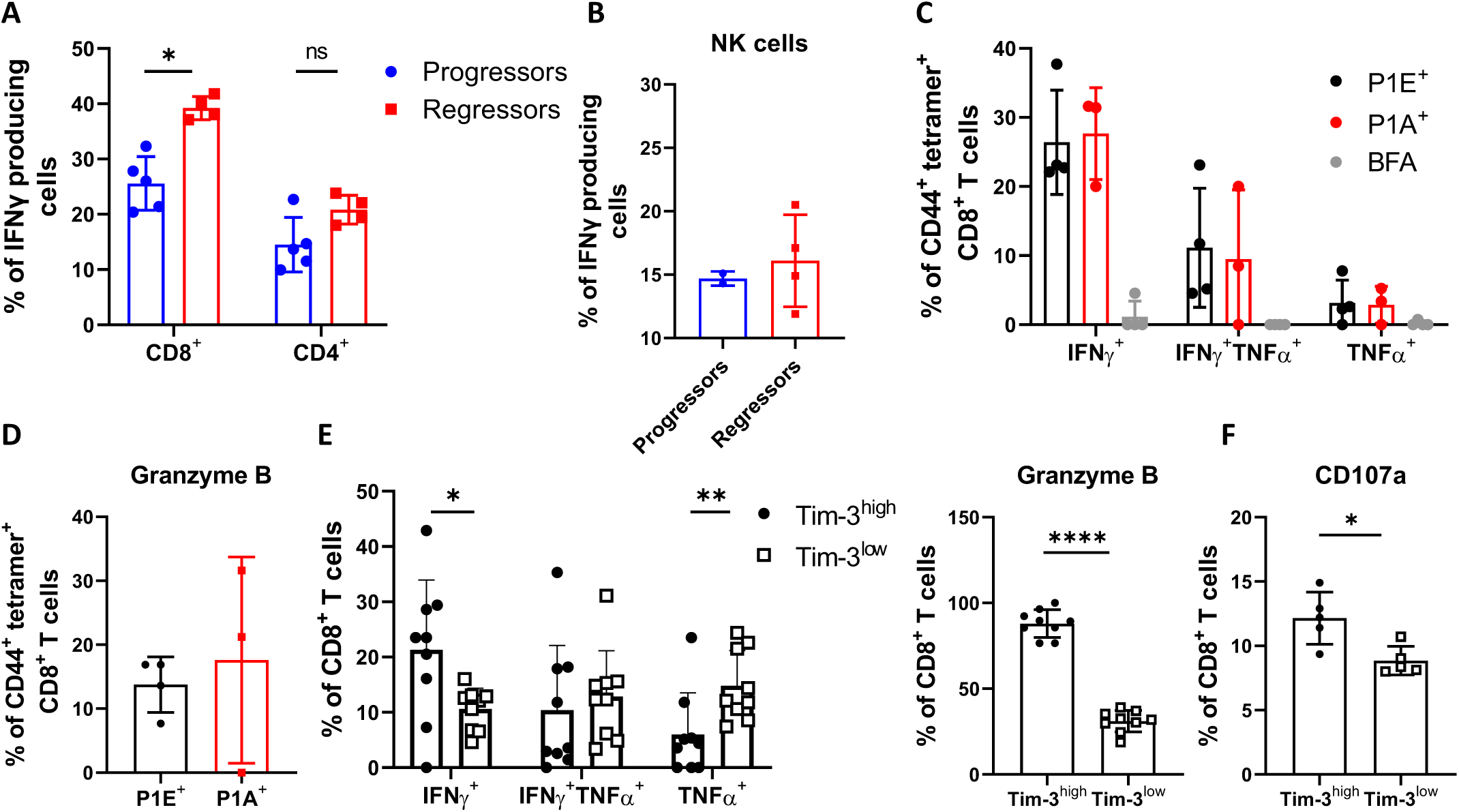
Identification of predominant IFNγ-producing cell population and their functional characterization. 1 x 10^6^ P815 mastocytoma cells were inoculated s.c. into both flanks of DBA/2 mice. Tumors from one flank were surgically extracted 10 days later and tumor-infiltrating immune cells analyzed by flow cytometry. Mice bearing contralateral tumors were treated with 250 µg of anti-PD-1 monoclonal antibodies from day 10 to day 13 and tumor growth was followed. Frequency of IFNγ producing cells among **A)** indicated T cell populations and **B)** NK cells. Tumor-infiltrating immune cells from one flank were harvested 11 days after tumor inoculation and analyzed by flow cytometry. Proportion of CD44^+^ P1E- and P1A-specific CD8^+^ T cells producing **C)** IFNγ, TNFα and **D)** granzyme B. **E)** Frequency of CD8^+^ T cells producing IFNγ, TNFα and granzyme B in Tim-3^high^ and Tim-3^low^ subsets. Production of IFNγ and TNFα was analyzed following stimulation with PMA/ionomycin. **F)** Proportion of Tim-3^high^ and Tim-3^low^ CD8^+^ T cells expressing CD107a. Data are representative of 1 experiment with 2-5 mice per group or E) show a pool of 2 independent experiments with 4-5 mice per experiment. Statistical significance was assessed by Mann-Whitney test (A, E, F) or unpaired t test (E). *p < 0.05, **p < 0.01, ***p < 0.001, ****p < 0.0001. The absence of asterisks in the graph indicates no significant difference between groups. BFA, brefeldin A.

**Figure S5.**
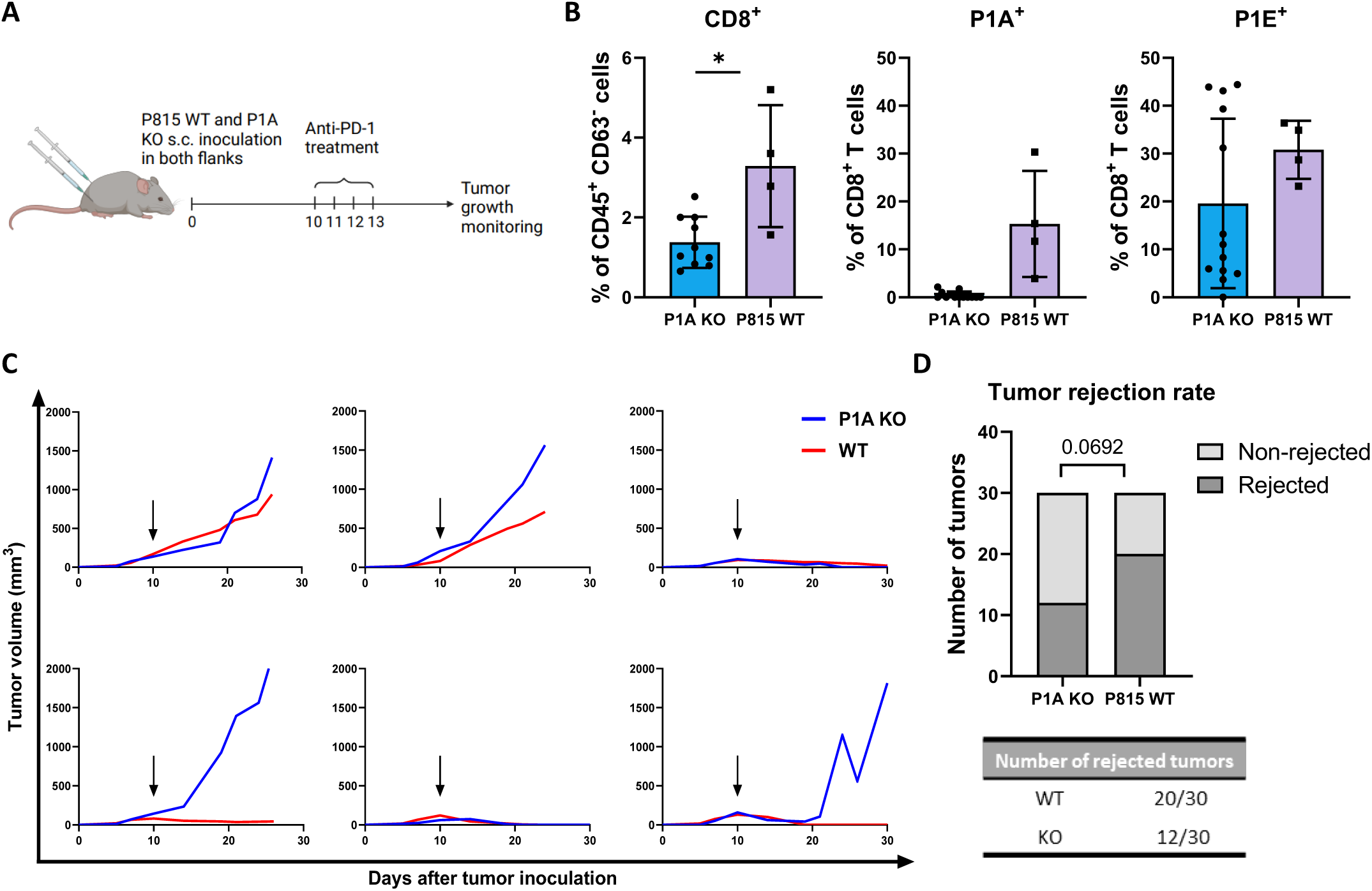
P1A-specific CD8^+^ T cell infiltration is not sufficient alone for a successful tumor response. DBA/2 mice were inoculated bilaterally with 1 x 10^6^ P815 mastocytoma and P1A KO cell lines respectively. Tumor-infiltrating immune cells were harvested 10 days later and analyzed by flow cytometry. **A)** Experimental set-up. Created with BioRender.com. **B)** Proportion of CD8^+^ T cells among CD45^+^ cells and proportion of P1A- and P1E-specific cells among CD8^+^ T cells in P1A KO and P815 WT tumors. DBA/2 mice were inoculated bilaterally with 1 x 10^6^ P815 mastocytoma cells in one flank and 1 x 10^6^ P1A KO cells in the contralateral flank. Tumor-bearing mice were treated with four daily injections of anti-PD-1 (Day 10 - 13) and tumor growth was monitored. **C)** Tumor growth of DBA/2 mice inoculated with P815 mastocytoma and P1A KO cells as indicated above. **D)** Tumor rejection rate of P815 WT and P1A KO tumor-bearing mice upon treatment with anti-PD-1. Data representative of B) 1 experiment with 4-13 mice per group and (C, D) 3 independent experiments with 8-13 mice per experiment. Statistical significance was assessed by Mann-Whitney test (B) and Fisher’s exact test (D). *p < 0.05.

**Figure S6.**
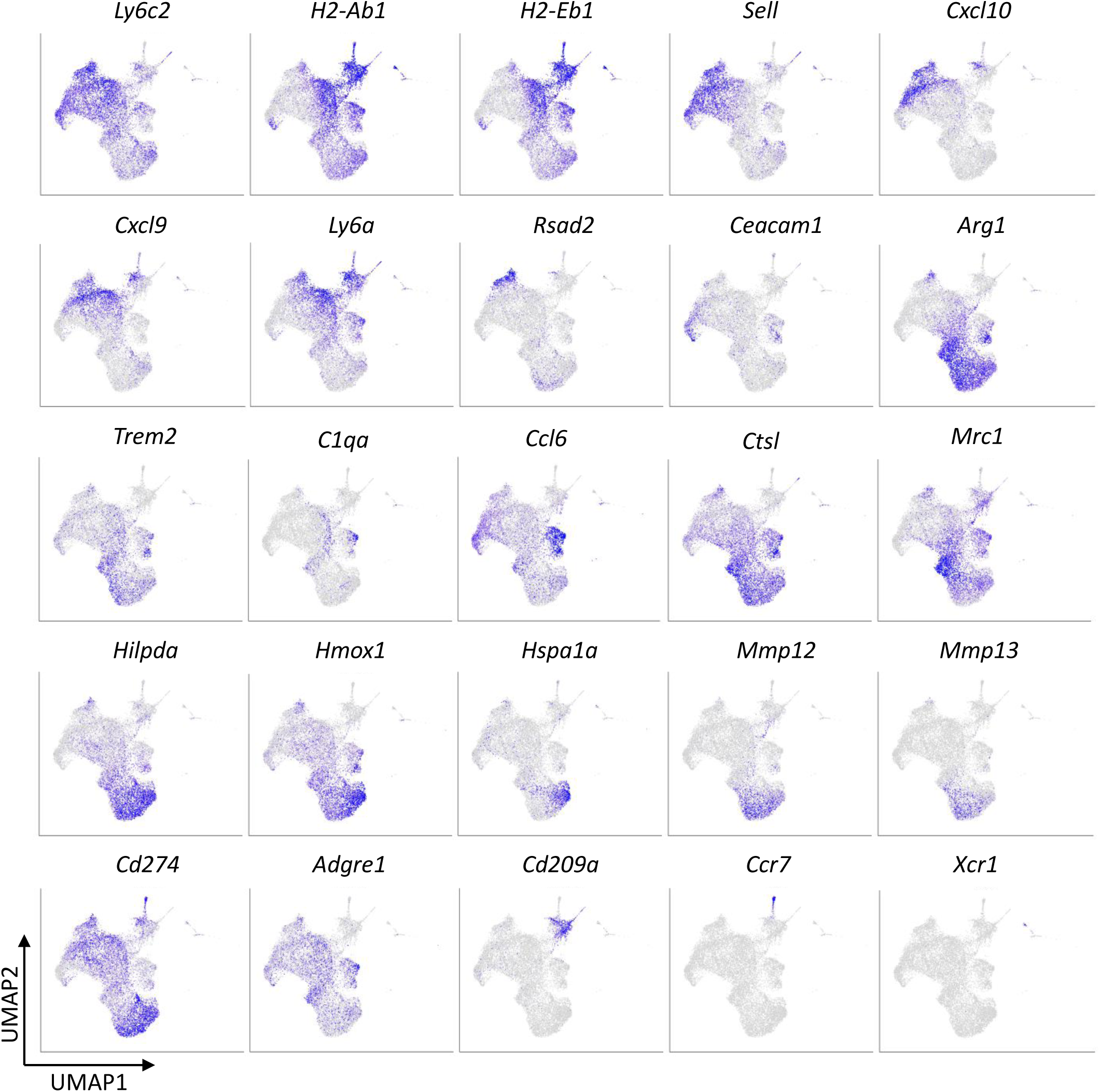
Transcriptional profiling of tumor infiltrating myeloid cells in pretreatment tumors. UMAP plots show the expression of markers used for the annotation of myeloid cell clusters.

**Figure S7.**
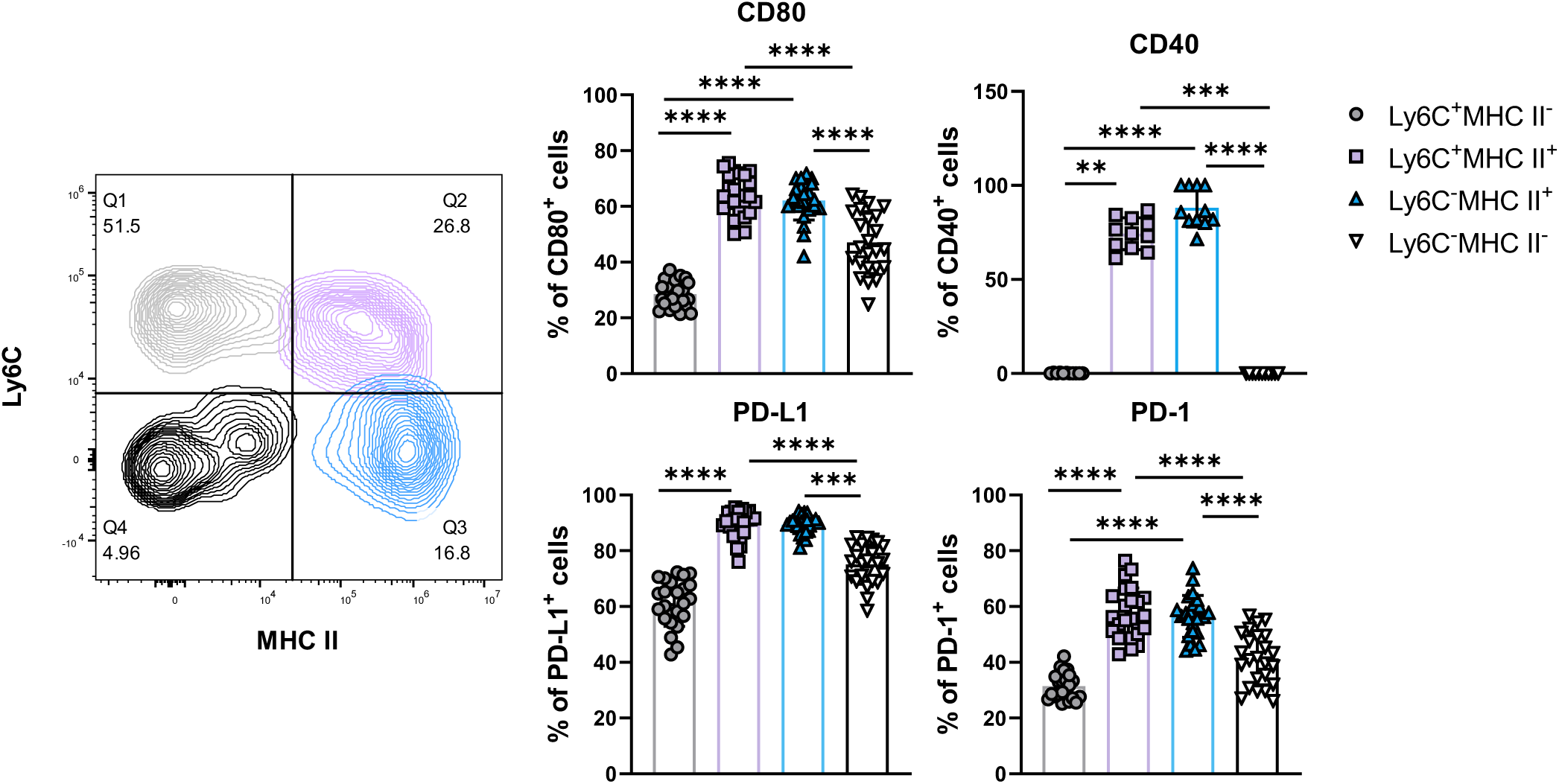
Enhanced antigen-presentation capacity is associated with subsets exhibiting elevated expression of MHC II. DBA/2 mice were inoculated s.c. with 1 x 10^6^ P815 mastocytoma cells in both flanks. Tumor infiltrating immune cells from the left flank were harvested 10 days after tumor inoculation and analyzed by flow cytometry. Representative FACS plot of Ly6C and MHC II showing four color-coded cell subsets. Frequency of CD80, CD40, PD-L1 and PD-1 - expressing cells within respective subsets. Data representative of 1 experiment with 12 mice (24 tumors in total) or 7 mice (11 tumors). Statistical significance was assessed by ordinary one-way ANOVA with Tukey’s multiple comparisons test or Kruskal-Wallis test with Dunn’s multiple comparisons test. **p < 0.01, ***p < 0.001, ****p < 0.0001.

**Figure S8.**
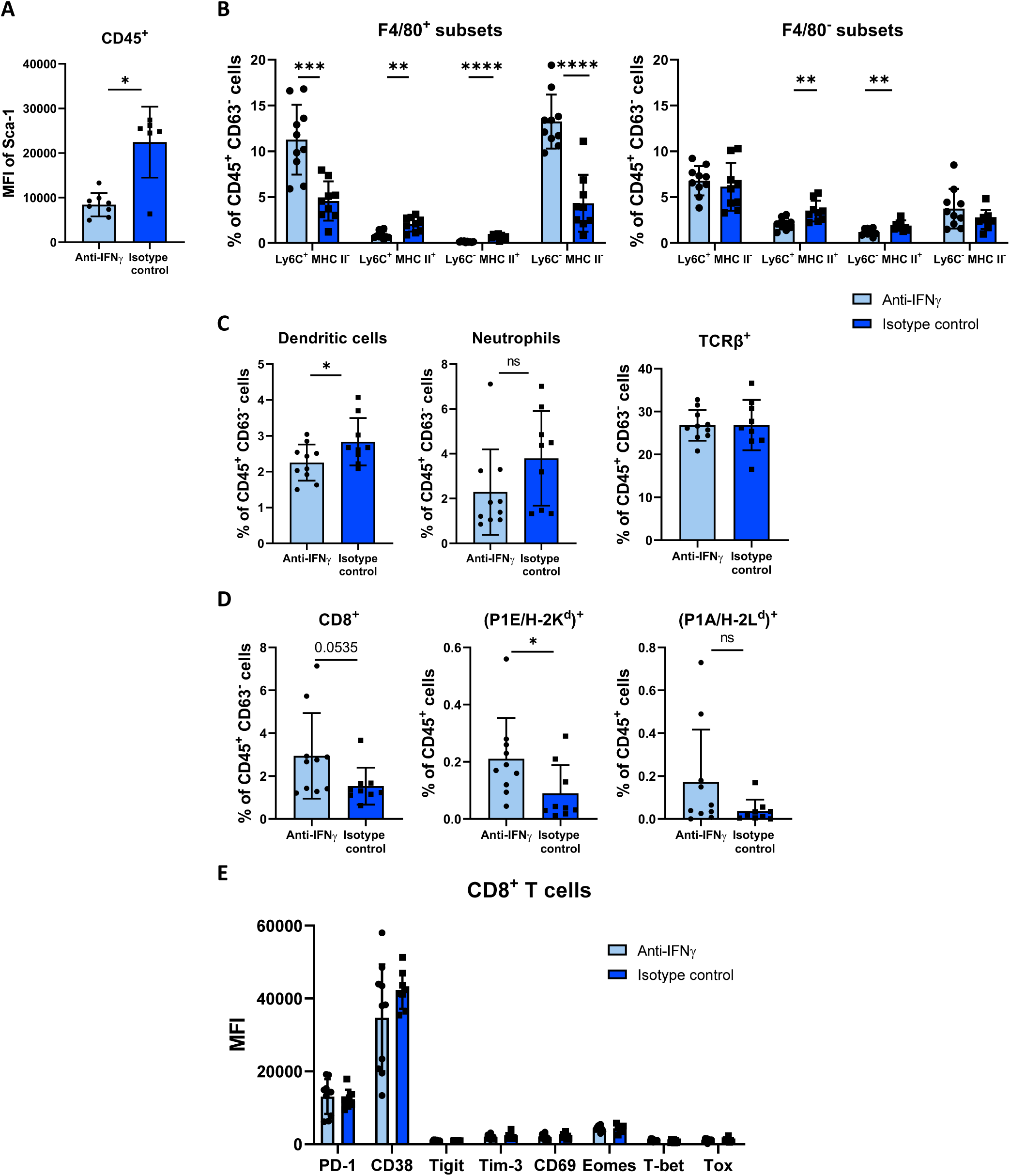
Effect of blocking IFNγ is particularly pronounced within myeloid cell populations. DBA/2 mice were inoculated s.c. with 2 x 10^6^ P815 cells in a single flank and treated intraperitonealy with 250 µg of anti-IFNγ monoclonal antibodies or isotype control. Tumors were harvested 10 days after tumor inoculation and analyzed by flow cytometry. **A)** MFI of Sca-1 in CD45^+^ cells. **B)** Proportion of F4/80^+^ and F4/80^-^ subsets among CD45^+^CD63^-^ population. **C)** Proportion of dendritic cells, neutrophils and TCRβ^+^ cells among CD45^+^CD63^-^. **D)** Frequency of total CD8^+^ T cells and P1E- and P1A-specific CD8^+^ T cells among CD45^+^ population. **E)** MFI of indicated markers and transcription factors in tumor-infiltrating CD8^+^ T cells. Data are representative of 1 experiment with 6-10 mice per group. Statistical significance was assessed by unpaired t test or Mann-Whitney test. *p < 0.05, **p < 0.01, ***p < 0.001, ****p < 0.0001. The absence of asterisks in the graph indicates no significant difference between groups.

## Notes

### Competing Interest Statement

The authors have declared no competing interest.

